# Density dependence on multiple spatial scales maintains spatial variation in both abundance and traits

**DOI:** 10.1101/759415

**Authors:** Koen J. van Benthem, Meike J. Wittmann

**Affiliations:** Department of Theoretical Biology, Bielefeld University, Bielefeld, Germany

**Keywords:** eco-evolutionary dynamics, population density, diversity, Allee effect, density-dependent selection

## Abstract

Population density affects fitness through various processes, such as mate finding and competition. The fitness of individuals in a population can in turn affect its density, making population density a key quantity linking ecological and evolutionary processes. Density effects are, however, rarely homogeneous. Different life-history processes can be affected by density over different spatial scales. In birds, for example, competition for food may depend on the number of birds nesting in the direct vicinity, while competition for nesting sites may occur over larger areas. Here we investigate how the effects of local density and of density in nearby patches can jointly affect the emergence of spatial variation in abundance as well as phenotypic diversification. We study a two-patch model that is described by coupled ordinary differential equations. The patches have no intrinsic differences: they both have the same fitness function that describes how an individual’s fitness depends on density in its own patch as well as the density in the other patch. We use a phase-space analysis, combined with a mathematical stability analysis to study the long-term behaviour of the system. Our results reveal that the mutual effect that the patches have on each other can lead to the emergence and long-term maintenance of a low and a high density patch. We then add traits and mutations to the model and show that different selection pressures in the high and low density patch can lead to diversification between these patches. Via eco-evolutionary feedbacks, this diversification can in turn lead to changes in the long-term population densities: under some parameter settings, both patches reach the same equilibrium density when mutations are absent, but different equilibrium densities when mutations are allowed. We thus show how, even in the absence of differences between patches, interactions between them can lead to differences in long-term population density, and potentially to trait diversification.

## 1. Introduction

Population density affects many aspects of an individual’s life, such as resource competition (Nicholson, 1957), parasite prevalence (Patterson and Ruckstuhl, 2013) and various aspects of the mating system, such as mate finding or competition for mating partners (Gascoigne et al., 2009; Kokko and Rankin, 2006). These processes can in turn affect lifetime reproductive success. For an individual it is thus advantageous to be adapted to the density it experiences. For example, at high density, investing in resource competition may pay off, whereas such an investment is futile when density is low. At low density, it may instead pay off more to invest in mate finding (Berec et al., 2018; Gascoigne et al., 2009). Such scenarios where the relative fitness of traits changes with density are referred to as density-dependent selection, a concept that has a long history (see MacArthur and Wilson, 1967). Although density-dependent selection is challenging to demonstrate (Travis et al., 2013), there are several clear examples. In a field population of great tits, fast exploratory behavior appears to be favored at low density and slow exploratory behavior at high density (Nicolaus et al., 2016). In experiments on *Drosophila* (Mueller, 1997; Mueller et al., 1991), populations were exposed to different densities, to which they adapted, most likely through evolution. Adaptation to density has also been demonstrated in moths (*Plodia interpunctella*), where males in an experiment adapted their reproductive strategy to the density experienced as larvae (Gage, 1995). Adaptation to density is also supported by observed patterns, such as the observed higher male aggressiveness in fig wasp species that tend to occur at smaller densities where killing another male yields the largest relative benefits (Reinhold, 2003).

An individual’s fitness can be affected by population density at more than one spatial scale, a phenomenon we call multi-scale density dependence. Multi-scale density depen dence should arise naturally if fitness is the result of multiple processes, e.g. occurring at different points in the life cycle. For example, birds may compete for high-quality nesting sites on a larger scale at the beginning of the breeding season, and then after settling on a nesting site compete for food more locally within their neighborhood (Rodenhouse et al., 2003). The effect of density may even be inverted depending on the spatial scale (Courchamp et al., 2008, box 2.7). For example, in arid vegetation, there is long-range competition for water, but also short-range faciliation because existing vegetation helps to retain water (Rietkerk, 2004). Similarly, mussels compete for food but may also benefit from a high local density, probably because it protects from waves (Gascoigne et al., 2005). In dogwood trees, when a focal patch is exposed to cicadas, the per capita number of attacks decreases with the tree density in that patch. However, whether cicadas decide to attack that patch, also depends on whether larger, more preferable, patches of trees are nearby (Cook et al., 2001). Individuals may thus be exposed to density effects at different scales simultaneously.

Here, we study how multi-scale density dependence affects spatial patterns of population density and variation in traits under density-dependent selection. We explore the possibility of obtaining a stable state with high-density patches that are being inhabited mostly by individuals that have a high density niche and low-density patches inhabited mostly by individuals adapted to low density. It has formerly been shown that spatial variation in density can emerge in homogeneous deterministic models, for example due to Allee effects (Gyllenberg and Hemminki, 1999), or due to the interplay of long-range competition either with small-scale facilitation (van de Koppel et al., 2005) or dispersal (Bolker and Pacala, 1997; Bolker, 2003; Sasaki, 1997). While these previous models have focused on ecological dynamics, we also include evolution of a trait under density-dependent selection.

The potential for adaptation to density is nontrivial because of the eco-evolutionary feedback loop (sometimes also referred to as eco-genetic feedback, Kokko and López-Sepulcre, 2007) that it is embedded in: while density may affect lifetime reproductive success, simultaneously changes in lifetime reproductive success also affect population density. The study of adaptation to spatial variation in density thus requires taking into account evolutionary and ecological processes simultaneously. So far, however, spatial models concerning these feedback loops have mainly focused on dispersal (Govaert et al., 2019). In our study, instead, we focus on the direct effect that patches can have on each other’s fitness.

We evaluate the capacity of multi-scale density dependent fitness to generate and maintain long-term differences in abundances between patches. Specifically, the fitness of an individual in our model is affected not only by the local density, but also by the density in a nearby patch. Such effects may emerge for example when a nearby patch attracts predators, that then spill over to the focal patch. Our model also includes the possibility of positive density dependence, which may occur for example when the nearby patch is attracting pollinators. By including traits into the model, we then study how subpopulations can adapt to their local density. For a plant population, for example, the investment into defenses against predators relative to the investment into attracting pollinators may be subject to density-dependent selection. However, simultaneously, the trait affects the density, thereby allowing for eco-evolutionary dynamics. We explore the conditions under which such a model can lead to diversification. Here we focus on allopatric diversification, that is, the evolution of different trait values in each patch.

## 2. Methods & Results

### 2.1. Model overview

We consider a population living in a habitat with two patches (Fig. 1). The patches may differ in the population density and in the trait distribution of the inhabiting individuals, but are otherwise identical. In particular, we assume for simplicity that they have the same area such that we can use density and population size or abundance interchangeably, but note that the results do not depend on this assumption. We first consider an ecological model where all individuals have the same trait value and there is no migration. We assume multi-scale density dependence in the sense that fitness in a patch depends not only on population density the patch itself, but also on the density in the other patch. Next, we consider an eco-evolutionary model where individuals differ in a trait under density-dependent selection and eco-evolutionary feedbacks between population density and trait distribution emerge. Finally, we evaluate whether the outcomes are robust to the inclusion of migration, stochasticity, and multi-locus genetics.

**Figure 1:**
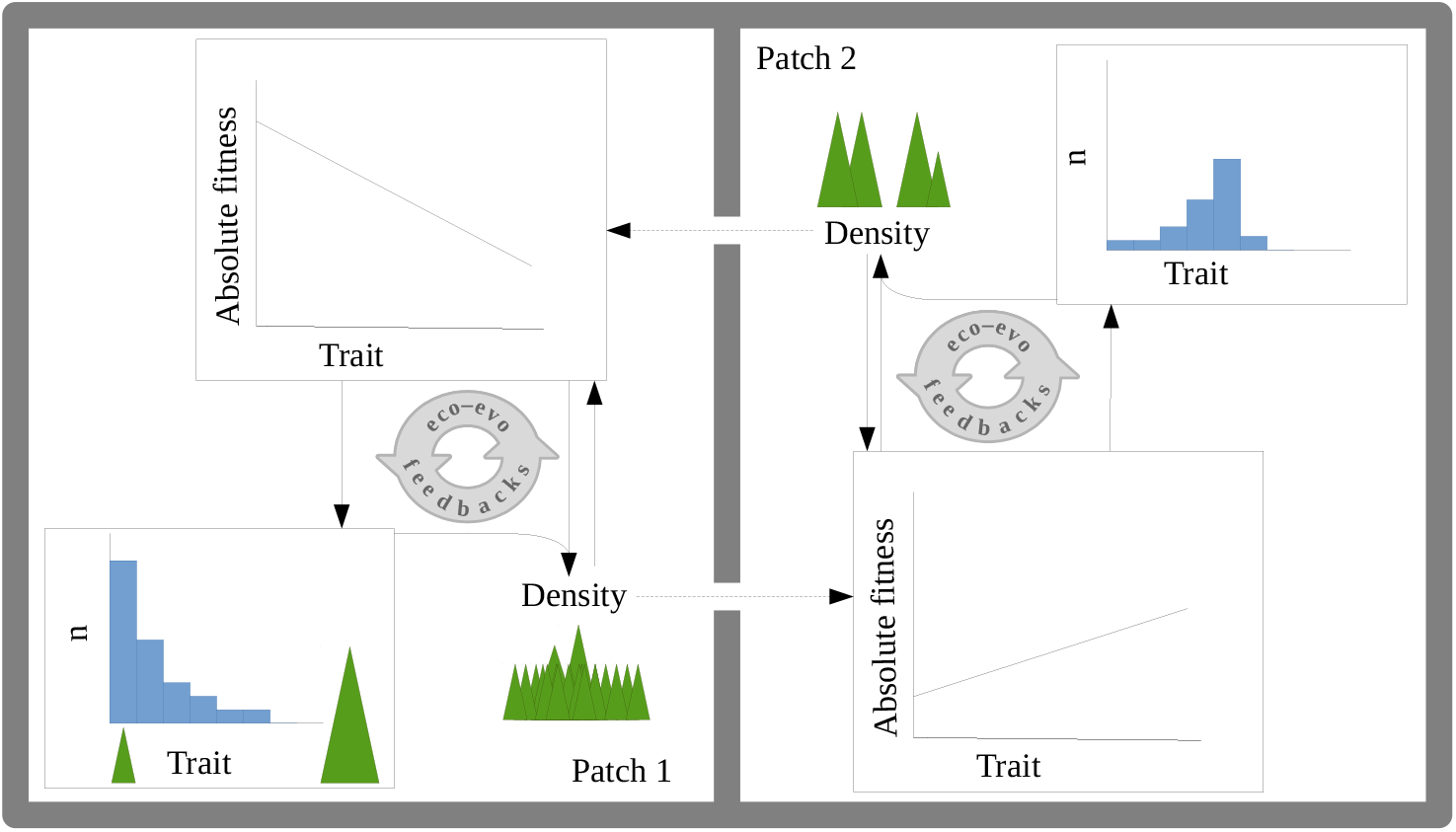
Overview of the ecological and evolutionary processes in our model. Absolute fitness differs between trait values and depends on density in the patch itself as well as on density in the other patch. Because of this density-dependent selection, the relationship between traits and fitness differs between patch 1 and 2. Absolute fitness then feeds back on population density but it also influences the evolution of the trait distribution (eco-evolutionary feedback).

### 2.2. Ecological model

The population dynamics in the two patches are described by two coupled differential equations:

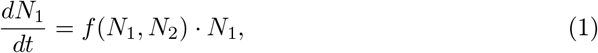

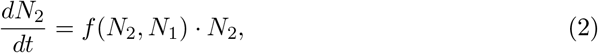

with *N*_*i*_ the density in patch *i* and *f* the per-capita growth rate or fitness function. Fitness depends on the abundances in both patches and a set of coefficients *c*_*α*_ (*α* ∈ [0, 1, 2, 3, 4]):

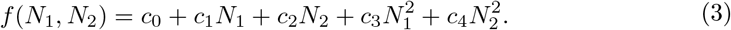

The coefficient *c*_0_ represents the intrinsic growth rate in an empty habitat. The linear coefficients *c*_1_ and *c*_2_ characterize the response to increasing density in the own and other patch, respectively, while these densities are still low. The quadratic coefficients *c*_3_ and *c*_4_ for response to the own and the other patch become increasingly important as densities increase and thus determine the high-density behaviour of the system. Individuals in this model experience density effects at two spatial scales. Density of their respective own patch influences fitness via the second and fourth term, and density in the respective other patch influences fitness via the third and fifth term. Both patches behave equally and exchanging their labels would not affect the results. Note that our model is mathematically speaking a special case of the model developed by Gerla and Mooij (2014) for the interaction between competing species in a single patch. With this different interpretation, their results are in line with parts of our results for the ecological model, as discussed below in more detail.

By choosing the parameters *c*_*α*_, our model can represent various scenarios. Here, we focus mainly on negative values for *c*_3_ and *c*_4_ to prevent populations from growing to infinity. When only considering the effect of the ‘own’ density on fitness, e.g. in patch 1, and keeping the density of the other patch fixed, 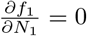 when 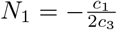. Since 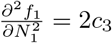, this point is a maximum if *c*_3_ < 0. If *c*_1_ is negative, the maximum is below zero and fitness decreases with *N*_1_ everywhere. If *c*_1_ is positive, the fitness maximum will be at a positive value of *N*_1_. Below the abundance at the extremum, per-capita fitness increases with increasing abundance. Hence, the system exhibits an Allee effect (Courchamp et al., 2008). At high densities, above the density at the extremum, the relation is inverted and percapita fitness decreases with increasing abundance, representing for example increasing resource competition or aggression. Depending also on the other parameters, the Allee effect with positive *c*_1_ might be strong with a negative per-capita growth rate at small densities or weak with a reduced but still positive per-capita growth rate at small densities (Courchamp et al., 2008). Similarly, when fixing the local density, fitness increases with the density in the other patch below 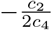 and decreases with density in the other patch above this value. Thus we can get either negative density dependence with respect to density in the other patch, or an analogue of a weak or strong Allee effect with respect to the density in the other patch. Of course all combinations of density-dependence scenarios with respect to the own and other patch are possible, thus accommodating many different biological situations.

#### 2.2.1. Equilibria

To determine the long-term spatial density patterns expected under this model, we compute the equilibria of the system (1) and (2) and assess their stability. From equation 1 it follows that the density in patch 1 is at equilibrium when *N*_1_ = 0 or *f* (*N*_1_*, N*_2_) = 0 or both, and analogously for patch 2. The overall system is at equilibrium when in each patch at least one of the conditions is met. Here, we focus on the cases where both patches have nonzero density. Hence they must both have a fitness equal to 0 and thus both of the following conditions need to be fulfilled:

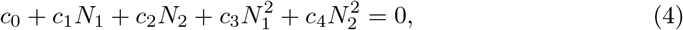

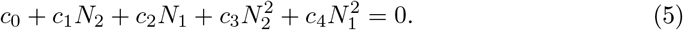

In the (*N*_1_,*N*_2_) phase plane, each of these equations corresponds to a conical section. Under our standard assumption that *c*_3_ and *c*_4_ are negative such that populations cannot grow to infinity, the isoclines are ellipses. The ellipse corresponding to the isocline of patch 1 is depicted in Fig. 2, including the equations for its center and its axes. Growth rates are positive in the interior of the ellipse and negative outside the ellipse.

**Figure 2:**
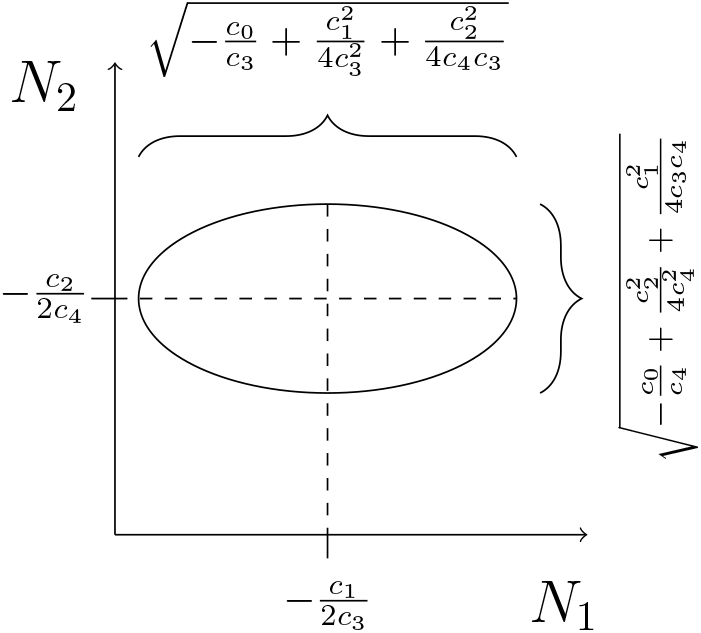
Geometric representation of the isocline. Isocline properties depend on ratios between coefficients only. Due to the isoclines containing no multiplication of *N*_1_ and *N*_2_, the axes of the ellipse are parallel to the *N*_1_- and *N*_2_-axis.

In general, two ellipses can have at most 4 intersections (Richter-Gebert, 2011), each of which is a possible equilibrium of the system (Fig. 3). Because the population dynamics in both patches are described by the same fitness function, the two corresponding isoclines are each other’s mirror image through the line *N*_1_ = *N*_2_. Hence, if the ellipses do not cross the diagonal line *N*_1_ = *N*_2_, they will not intersect and hence, there will be no overall equilibrium, except for the one in which both patches are empty (Fig. 3 a). If one ellipse crosses the line *N*_1_ = *N*_2_ in two points, due to the symmetry of the system, the other ellipse will also intersect the diagonal and thereby the first ellipse at these points (Fig. 3 b). On the diagonal there are thus either 0 or 2 intersections. However, whether these intersections are meaningful depends on whether they lie in the positive quadrant, since negative values for abundance are not biologically relevant. Hence, in addition to the origin where both patches are empty, there can be 0, 1 or 2 biologically meaningful equilibria on the diagonal. These equilibria are all of the type where both patches have the same abundance. If the ellipses cross the diagonal, they can additionally intersect in two, and only two, additional points away from the diagonal. Due to the symmetry of the system, if a single intersection away from the diagonal exists, so must its mirror image through the line *N*_1_ = *N*_2_. These cases, with 4 intersections, allow for the system to be in an equilibrium with both patches having different abundances, even though they are being governed by the same fitness function (Fig. 3 c).

**Figure 3:**
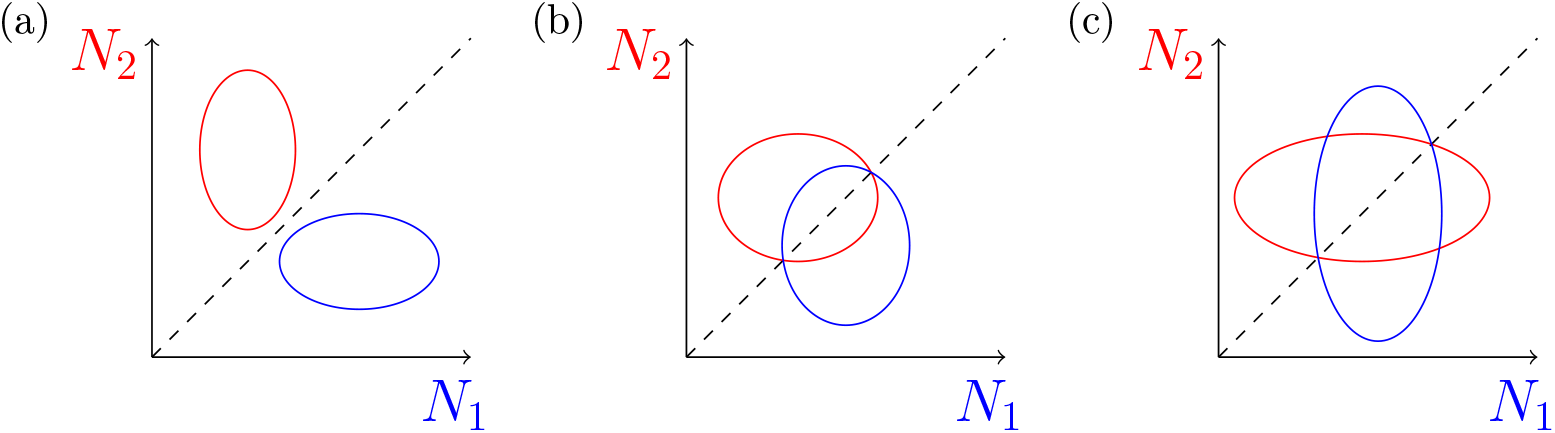
Possible number of intersections for two mirroring ellipses. This figure only illustrates the isoclines where *f_i_* = 0, the isoclines for *N*_*i*_ = 0 correspond to the x and y-axes and the ellipses can also intersect these.

The values of the equilibria can be obtained by solving equations 4 and 5 simultaneously.

First, we subtract the two equations from each other and obtain:

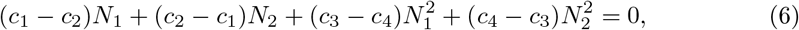

which can be rewritten as:

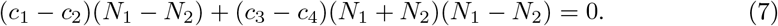

This can be factorized into a term depending on the difference between the abundances and a term depending on the total abundance:

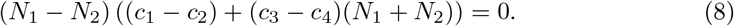

This condition is fulfilled when either of the two factors is zero, that is if *N*_1_ = *N*_2_ or

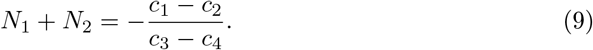

In the special case when *c*_3_ = *c*_4_, the second equilibrium does not exist. Here, we are most interested in the second solution because it allows for equilibria where the two patches contain a different density. We can now use this solution to eliminate *N*_2_ from equations 4 by setting 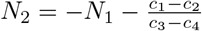 and regrouping:

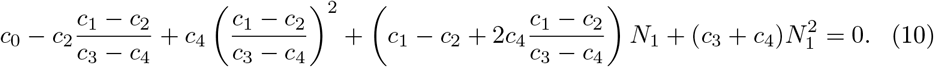

We will solve this equation using the quadratic formula. Before we do so, we can slightly simplify the higher order terms in the above equation by multiplying all terms with 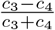:

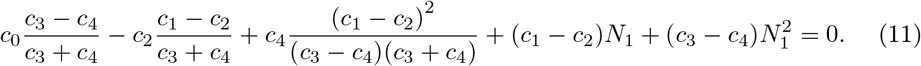

This can now be solved using the quadratic equation:

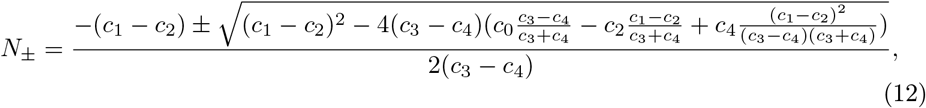

which can be written as:

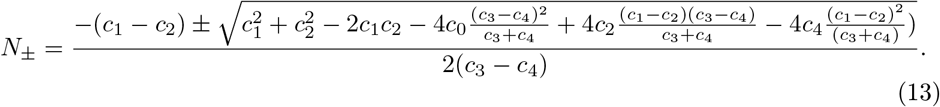

Some rearrangement of these terms allows this expression to be written as:

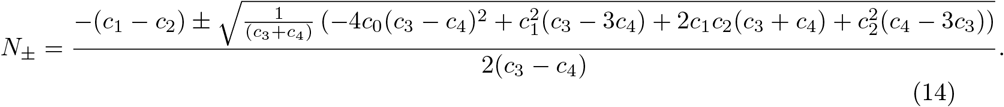

Since *N*_+_ and *N*_−_ represent densities, they should both be positive. A necessary condition to achieve this is for 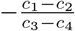 to be positive. This means that either *c*_1_ > *c*_2_ and *c*_3_ < *c*_4_ or *c*_1_ < *c*_2_ and *c*_3_ > *c*_4_. In most systems, *c*_3_ and *c*_4_ will be negative, to prevent explosive population growth. For these systems, this necessary condition can be written as *c*_1_ > *c*_2_ and |*c*_3_| > |*c*_4_| or *c*_1_ < *c*_2_ and |*c*_3_| < |*c*_4_|. If *c*_1_ and *c*_2_ are both positive, the two equilibria can thus only be meaningful when the stronger linear response in the fitness function is coupled to the stronger quadratic response; that is, the positive density dependence at low density and the negative density dependence at high density should both be stronger for the own patch or both be stronger for the other patch.

Note that the equilibria are constant under scaling: when all coefficients are multiplied with the same positive constant, the equilibria are unaltered. This is in agreement with the fact that the ellipses only depend on ratios between coefficients and not on their absolute values (see Fig. 2).

#### 2.2.2. Stability analysis

The stability of the equilibria can be obtained through the Jacobian matrix

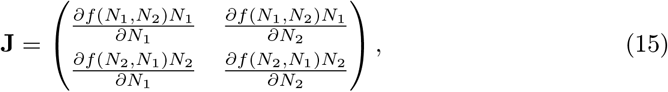

and hence

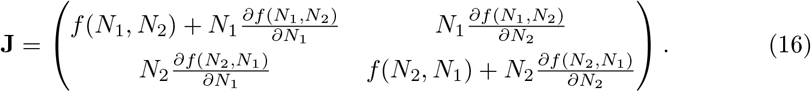

At the nontrivial equilibrium, *f*(*N*_1_, *N*_2_) = *f*(*N*_2_, *N*_1_) = 0 and thus:

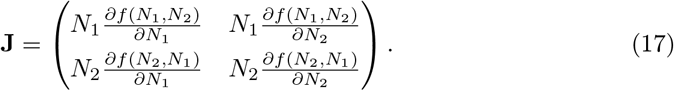

If both eigenvalues of this matrix, evaluated at an equilibrium of interest, have a negative real part, the equilibrium is stable. Using the quadratic formula to solve the corresponding characteristic equation, we find the two eigenvalues:

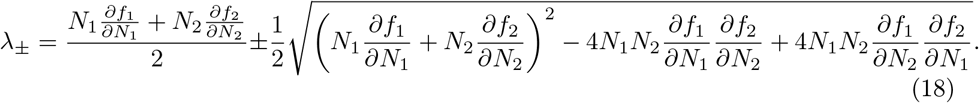

Here, we used the short-hand notation *f*_1_ = *f* (*N*_1_*, N*_2_) and *f*_2_ = *f* (*N*_2_*, N*_1_). After rear-ranging, we finally obtain

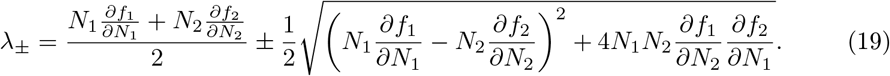

To assess stability, it suffices to check the eigenvalue with the largest real part. Since the real part of *λ*_+_ is greater or equal to the real part of *λ*_−_, the condition for stability thus becomes:

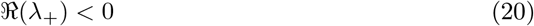

A necessary but not sufficient condition for this is:

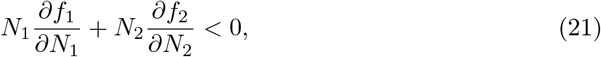

or:

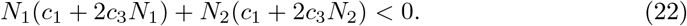

We use equation 20 to evaluate the stability of the equilibria from equation (14) by setting *N*_1_ = *N*_+_ and *N*_2_ = *N*_−_ (or vice versa). As noted above, these values remain the same as long as all coefficients keep their relative values and sign. Furthermore, if the fitness function is multiplied by a positive, real constant, the eigenvalues of the Jacobian matrix also scale with this constant and hence the stability does not change (multiplication by a positive number does not affect the sign). Hence a temporally varying environmental factor that acts by scaling the coefficients and hence the fitness function uniformly in both patches, would not alter the equilibria nor their stability. Such environmental variation may however change the basin of attraction of equilibria.

#### 2.2.3. Ecological model results

Fig. 4 (top panels) shows an example system where at equilibrium both patches can have different densities. The left panel shows abundance time series, while the corresponding phase space trajectories are shown on the right. The colors refer to initial conditions (same color means same initial conditions). From the right panel it becomes clear that all trajectories end on an intersection between two isoclines. However, the specific equilibrium that the system reaches depends on its initial values for *N*_1_ and *N*_2_. The coefficients in the bottom panels were equal to those in the top panels, but with the effects of ‘own’ and ‘other’ patch exchanged (*c*_1_ ↔ *c*_2_ and *c*_3_ ↔ *c*_4_). In this system, the equilibria where the patches contain different nonzero abundances are unstable. Instead, one of the equilibria with equal abundance in both patches is stable. Furthermore, additional equilibria, where one of the two patches goes extinct, have become stable.

**Figure 4:**
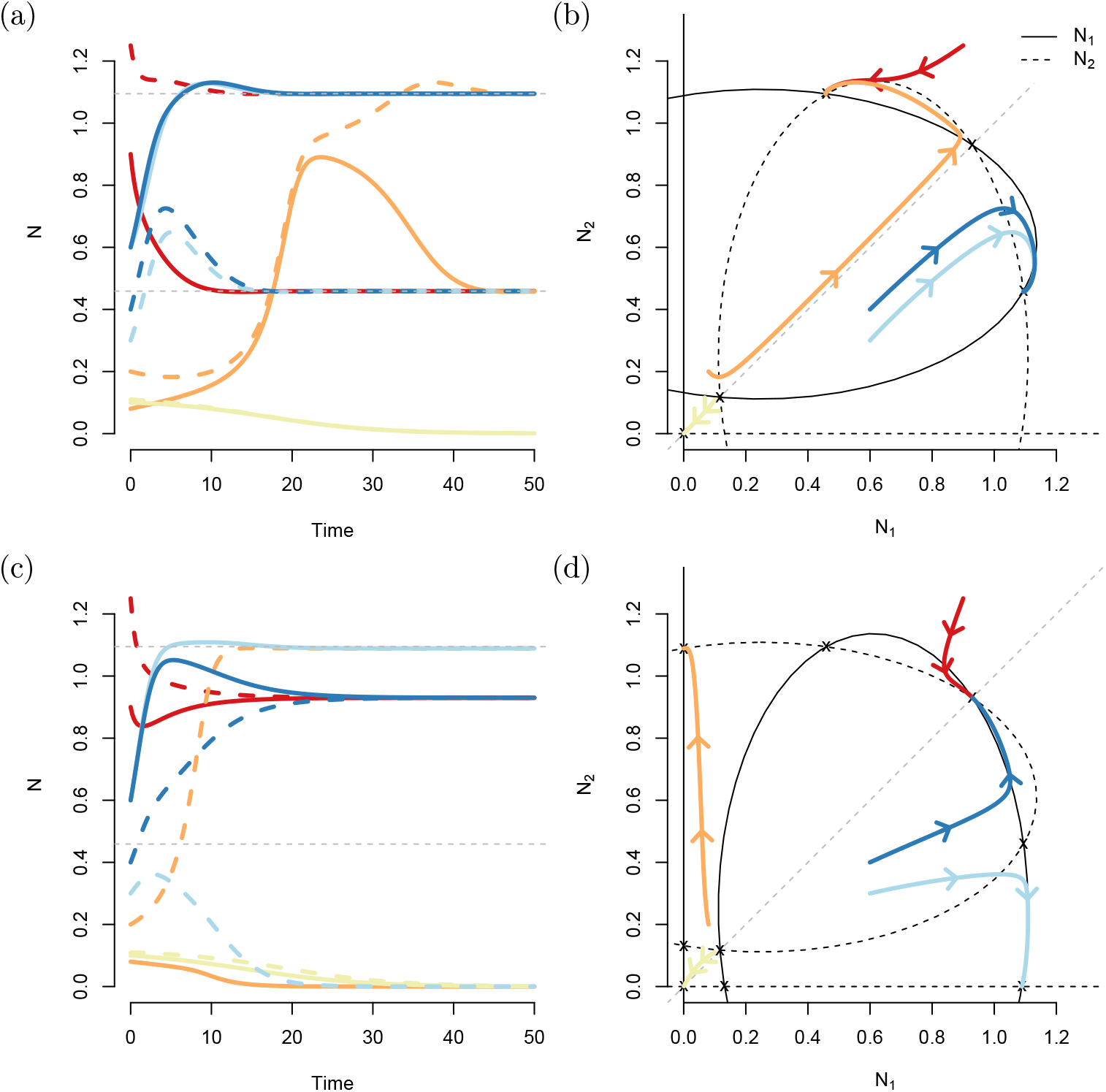
Example time series (a,c) and corresponding trajectories and isoclines in phase space (c,d). Different colors correspond to different initial values. The parameter values used for (a,b) were: *c*_0_ = −0.148, *c*_1_ = 0.162, *c*_2_ = 1.262, *c*_3_ = −0.326, and *c*_4_ = −1.034. The parameters for (c,d) were *c*_0_ = −0.148, *c*_1_ = 1.262, *c*_2_ = 0.162, *c*_3_ = −1.034, and *c*_4_ = −0.326. The bottom panels thus describe a system in which the effects of the ‘own’ patch and the ‘other’ patch have been exchanged. The ellipses also exchange identity, although their shape and intersection remain the same. The stability of the equilibria did change however.

The values of the coefficients determine which equilibria exist and which of these are stable. With equations 14, 19 and 20, it is possible to calculate the equilibria for any set of coefficients and evaluate their stability. The full parameter space is five-dimensional, but we evaluated the equilibria and their stability only at two-dimensional cross sections of that space (Fig. 5). Each cross section describes the effect of two of the coefficients, whilst keeping the remaining three coefficients at their value from the top panels of Fig. 4. The figure shows that most parameter combinations do not lead to an equilibrium in which both patches contain a different number of individuals. However, the region within which both patches may settle to different abundances is non-negligible. Small changes in the coefficients around the values from Fig. 4 are therefore not expected to lead to qualitative differences in the outcomes.

**Figure 5:**
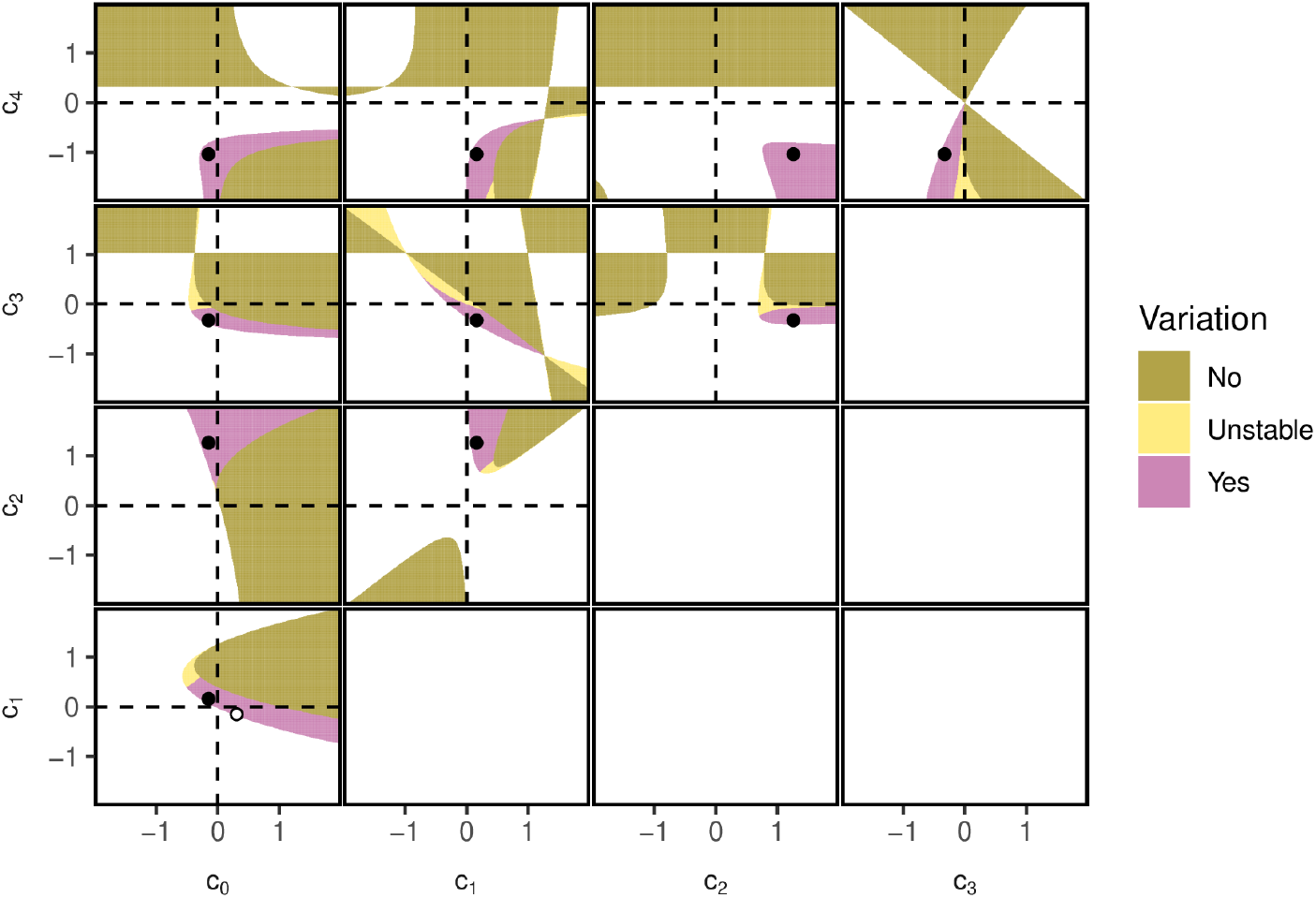
Regions in parameter space where density variation in space can be stable. Only the upper diagonal graphs are shown. Here, in the white regions, equation 14 returned equilibria with a nonzero imaginary part. In the ‘No’ region, the obtained equilibria were real, but either unreachable (at least one of them was negative) or the equilibria for *N*_1_ and *N*_2_ were the same. Finally, there were cases where the two equilibria were real, positive, and different. These cases were again subdivided in cases where the equilibrium was stable (‘Yes’) and where it was not (‘Unstable’). The black dots correspond to the parameter settings that were used in the top panels of Fig. 4. The white dot in the *c*_0_-*c*_1_ panel corresponds to the parameter setting of *c*_0_ and *c*_1_ in Fig. 6.

**Figure 6:**
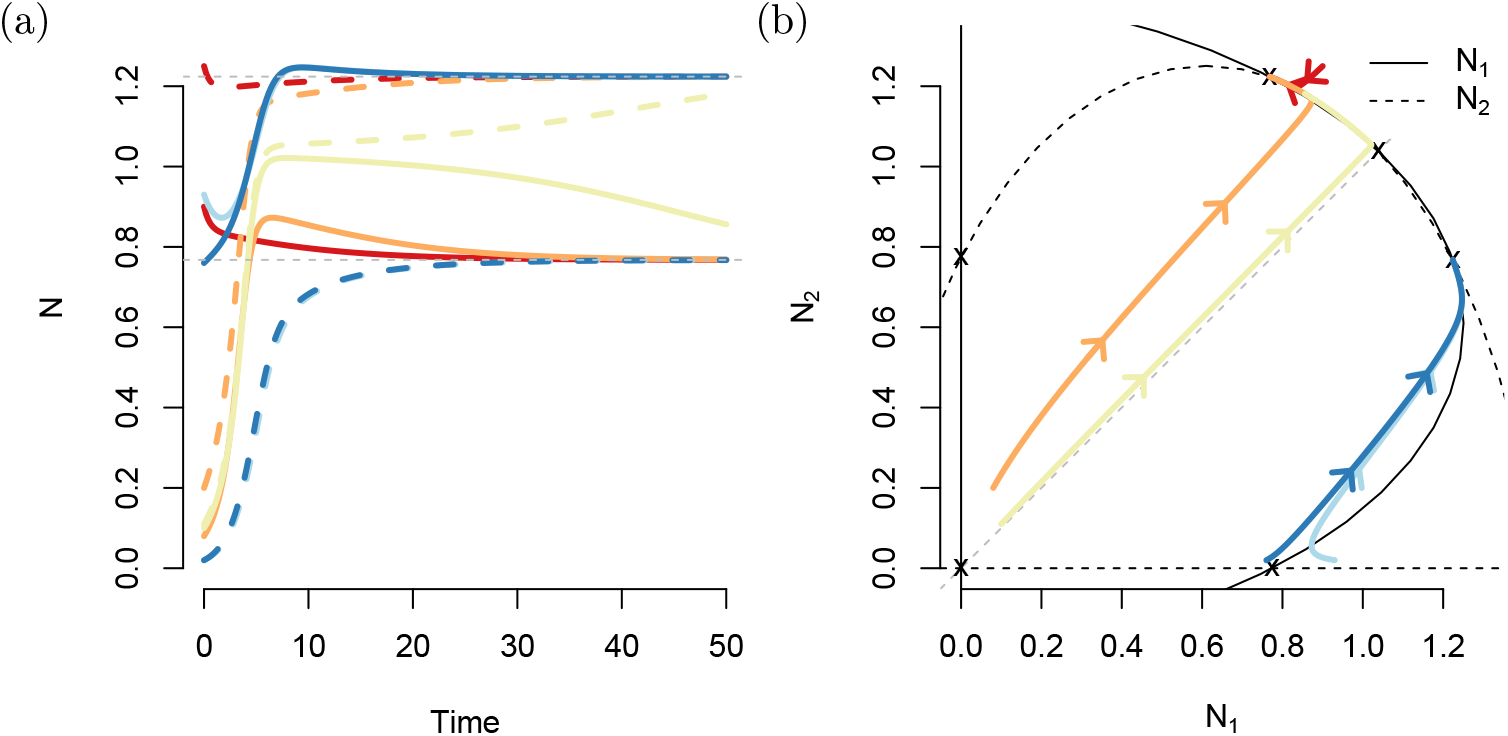
Example time series (a) and corresponding trajectories and isoclines in phase space (b). Different colors correspond to different initial values. The parameter values were: *c*_0_ = 0.31, *c*_1_ = −0.148, *c*_2_ = 1.262, *c*_3_ = −0.326, and *c*_4_ = −1.034.

Above, we remarked that a necessary condition for the existence of meaningful asymmetric equilibria is that the strength of the response to the other patch is stronger than the response to the own patch in both linear and quadratic term, or weaker in both linear and quadratic term. Here we have explored the parameter space around a point where the other patch has a stronger effect and thus we observe stable variation in abundance when *c*_2_ > *c*1 (see 3rd row, 2nd column in Fig. 5) and when |*c*_4_| > |*c*_3_| (4th row, 4th column in Fig. 5). Exploration of the *c*_1_-*c*_0_ parameter space (4^th^ row, 1^st^ column in Fig. 5) reveals that stable spatial density variation should be possible also for cases where increasing density in the own patch has a negative effect even at low density (*c*_1_ < 0). Time series and isoclines for a parameter combination in this region are shown in Fig. 6. In this example, both *c*_1_ and *c*_3_ are negative, meaning that there is no Allee effect acting directly within the focal patch, although *c*_2_ is still positive, leading to positive effects of density in the other patch on the fitness in the focal patch at low density in the other patch. Furthermore, all trajectories in this example converge to the asymmetric equilibria, which turn out to be the only stable equilibria.

### 2.3. Eco-evolutionary model

Long-term differences in population density may lead to diversification in traits under density-dependent selection, which may in turn affect the densities. In order to allow for such eco-evolutionary feedbacks, we now include a trait, *z*, that takes values between 0 and 1. The trait affects the fitness through two additional terms: the first quantifies a density-independent effect of the trait value on the fitness (*c*_5_*z*), while the second describes a density-dependent effect of the trait value (*c*_6_*zN_i_*):

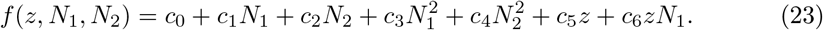

Examples of traits whose fitness consequences are affected by density are investment in attributes for fighting or pheromone production for mate finding. Note that in this model, the selection on *z* changes with the density in the own patch, but not with the density in the other patch.

Now, not only the population size, but also the trait distribution matters. We track the trait distribution by dividing the trait space into 100 discrete bins, with *z_b_* the trait value of bin *b* and all the *z_b_* evenly spaced between 0 and 1. The total abundance in patch 1 is simply the sum of the number of individuals in all size classes *b* in the patch:

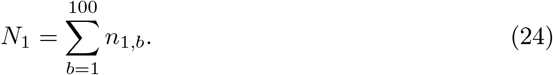

The abundances *n*_1*,b*_ change through reproduction, as described by the fitness function, as well as through mutations. We treat mutations deterministically, such that individuals in bin *b* mutates away to either bin *b −* 1 or *b* + 1 at a rate *μ*. In the first and the last bin, the trait value can only move in one direction, and accordingly, the mutation rate in this bin is halved 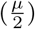. The dynamics of this system are described by a set of differential equations:

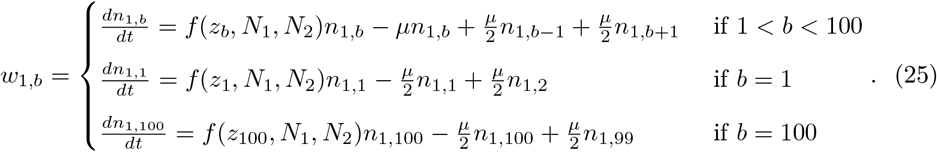

The equations for the second patch are analogous. This yields 200 coupled differential equations, that we initially solved numerically using the package deSolve in R (R Core Team, 2018; Soetaert et al., 2010).

#### 2.3.1. Equilibria

The numerically obtained equilibria were then compared to a direct calculation of the equilibria when *μ →* 0. For a given combination of *N*_1_, *c*_5_, and *c*_6_, the fitness function is monotonic in *z*. If *c*_5_ + *c*_6_*N*_1_ > 0, larger trait values will be selected for (smaller trait values if *c*_5_ + *c*_6_*N*_1_ < 0). We therefore hypothesize the average trait value of a patch to end up at either boundary (0 or 1). Since population density and thus the direction of selection can differ between patches, the two-patch system can potentially have four equilibria. We solved for the equilibria by solving the following four sets of two equations:

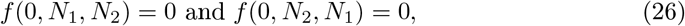

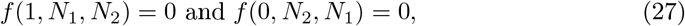

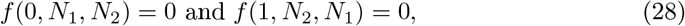

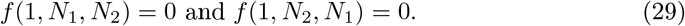

This procedure is equivalent to intersecting the two ellipses with *z* = 0 and *z* = 1 for the first patch with the two analogous ellipses of the other patch in all 2 × 2 combinations. The exception occurs when *c*_5_ + *c*_6_*N*_*i*_ = 0; in this case there is no selection on the trait value and hence, the equilibrium has become independent of the trait value, and we should still find it when *z* = 0 or *z* = 1. We used Mathematica to find the solutions for equations (26) – (29).

#### 2.3.2. Stability

Furthermore, we evaluated the stability, by calculating the dominant eigenvalue of the Jacobian matrix for the system of 200 differential equations. At each equilibrium of the system, we compute the Jacobian

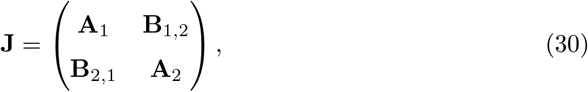

with:

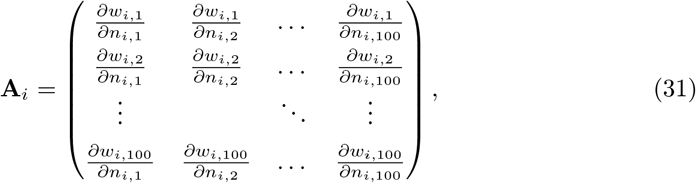

and

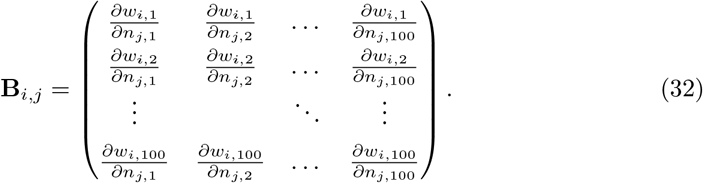

When estimating the actual stability of an equilibrium, we set the mutation rate to 0. The reason is, that with a nonzero mutation rate, the final trait distribution will not be monomorphic at either *z* = 0 or *z* = 1, due to the selection-mutation balance. However, this slight mismatch may also affect the value of the equilibria. To circumvent this issue, we evaluate the stability in the limit *μ* → 0, where the mutation-selection balance is also expected to be fully favoring selection. Finally, we confirmed the stability metrics using numerical solutions to the system of differential equations, as presented in SI S1.

#### 2.3.3. Eco-evolutionary model results

Fig. 7 shows time series generated by the model with traits included. Initially, the populations were monomorphic, with all individuals having a trait value of either *z* = 0 or *z* = 1. When the initial trait value in the population was *z* = 0, both patches reached very similar equilibrium population density, regardless of mutation rate (red and blue lines). Hence, the system allows for stable spatial density variation even without variation in trait value. In contrast, when starting with a monomorphic population with trait value *z* = 1, the presence of mutations qualitatively affected the population dynamics (orange and yellow lines). In the absence of mutations, both patches reached the same density (orange line). With mutations, however, the system reached a final equilibrium in which both patches contained a different number of individuals, as well as different trait distributions (yellow lines). In this case the system was thus governed by eco-evolutionary feedbacks.

**Figure 7:**
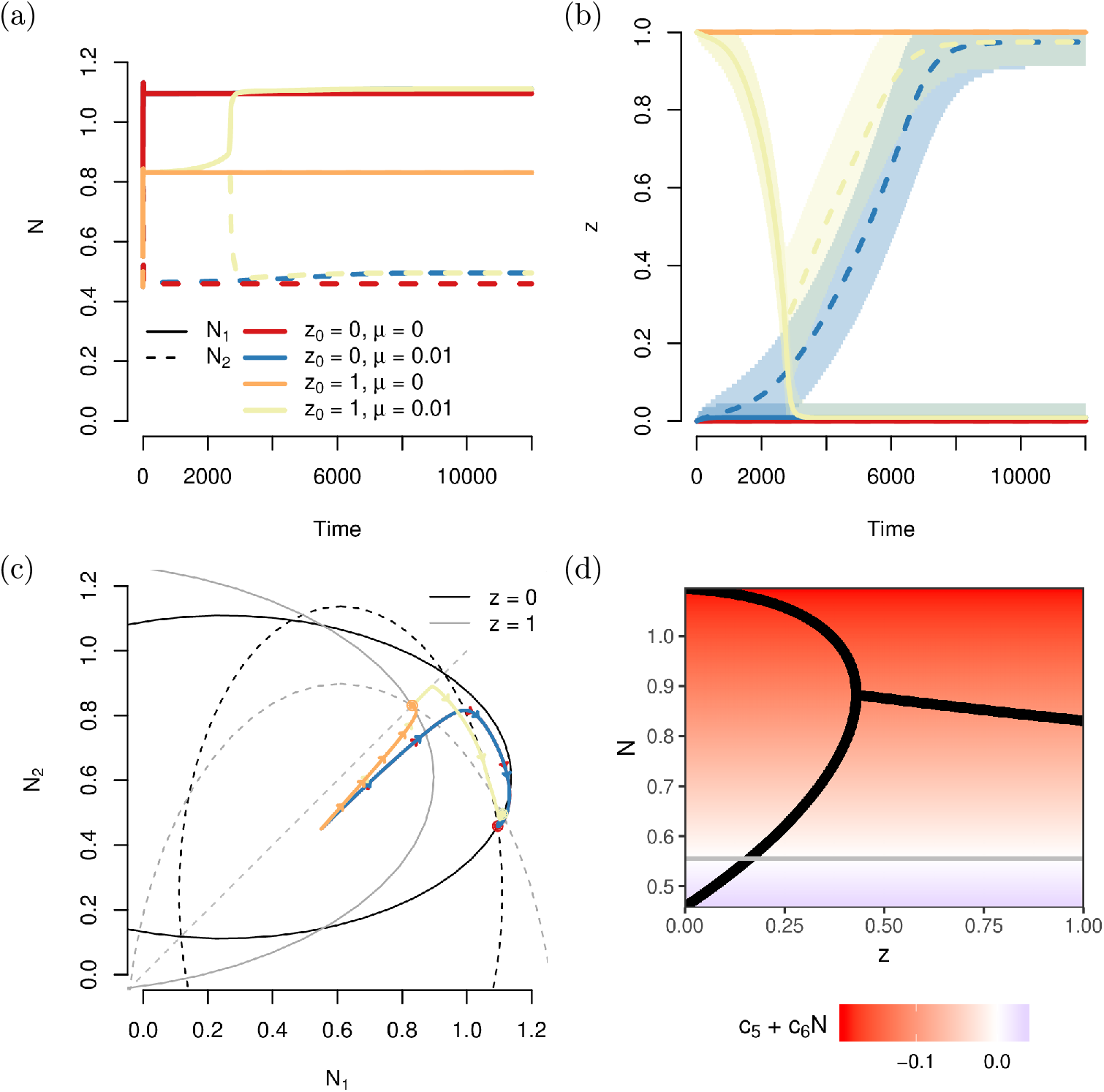
Example time series for the eco-evolutionary model. Different colors represent different model runs where mutations were either present (*μ* = 0.01) or not (*μ* = 0) and that started with a monomorphic population with starting trait value *z*_0_ either 0 or 1. (a) Abundance time series. The red and blue trajectories largely overlap for the first 2000 time steps, as do the yellow and orange lines. (b) Corresponding mean trait values and the spread, shown as the regions in trait space that contained 95% of the individuals of each patch. For the scenarios without mutations (orange and red line), the trait values in both patches completely overlap. For the scenario with mutations starting at *z* = 1 (the yellow lines), initially the lines in both patches overlap, but around time 2500, the trait values in the two patches start to diverge. (c) Trajectories in phase space. Also drawn are the isoclines at *z* = 0 (black) and *z* = 1 (grey). (d) Equilibrium densities for monomorphic populations with trait value *z*. On the left side of the graph, at any given value of *z* two branches exist, indicating the two different densities that the two patches will tend to. In the background, the direction of selection at any given density is shown, with red values referring to selection for smaller trait values and blue colors to selection for larger trait values. The grey line corresponds to the density at which selection vanishes. Parameter values: *c*_0_ = −0.148, *c*_1_ = 0.162, *c*_2_ = 1.262, *c*_3_ = −0.326, *c*_4_ = −1.034, *c*_5_ = 0.194 and *c*_6_ = −0.3492.

In the region of parameter space explored here, simultaneous maintenance of variation in abundance and diversification of trait values depends strongly on the values of the parameters *c*_0_ to *c*_6_. In Fig. 8, we again varied two parameters at a time while keeping the others constant at the values in Fig. 7. We divided the parameter space into regions with at least one stable equilibrium with variation in both *z* and *N*, regions with no stable joint variability but with the possibility of stable variation in *N*. We also looked for regions with the possibility of stable variation just in *z* but did not find any.

**Figure 8:**
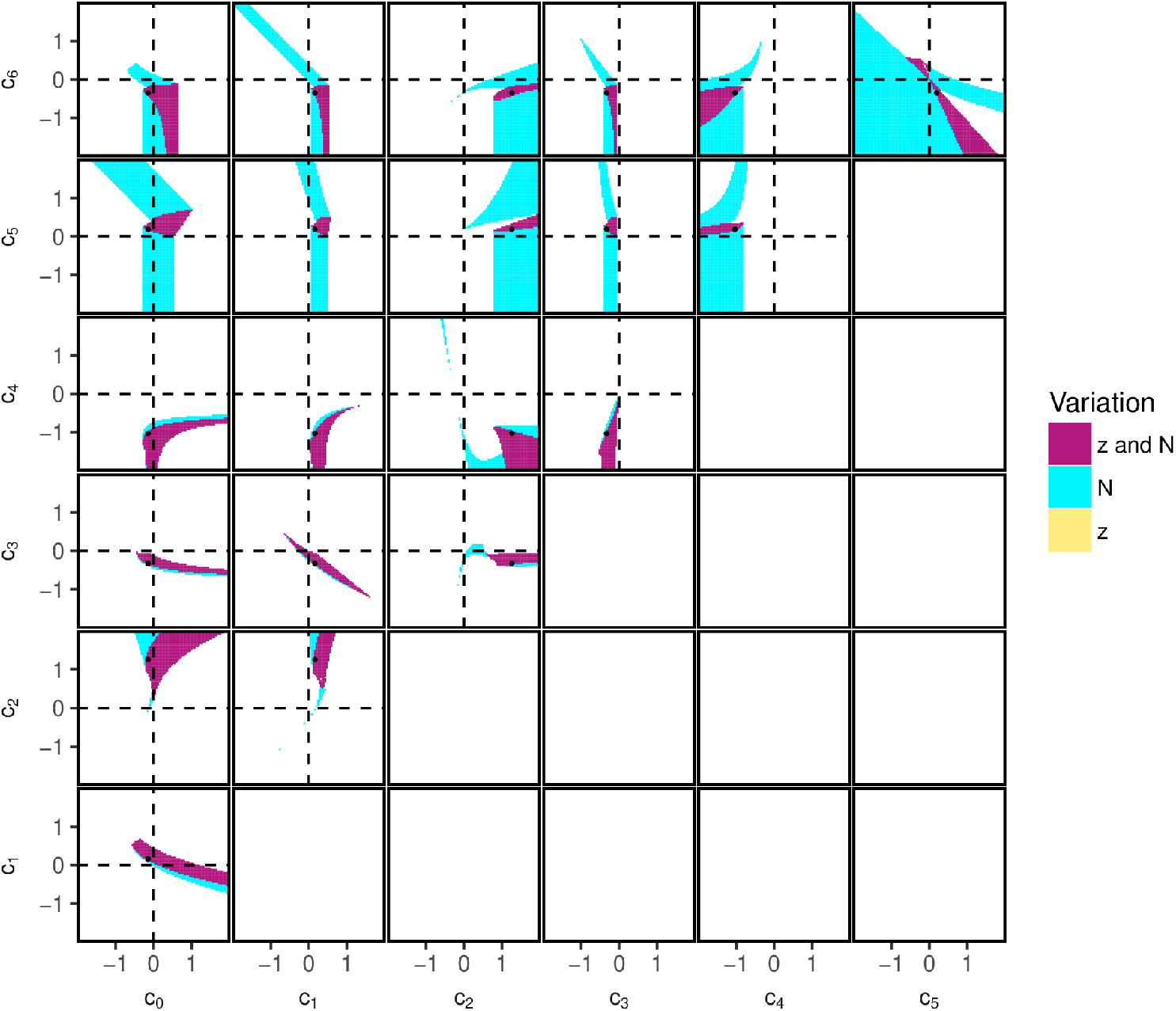
Regions in parameter space where spatial variation in abundance and diversification of trait values can occur. The shown regions are based on the properties of the asymmetric, equilibria obtained by intersecting the ellipses with trait value *z* = 0 and *z* = 1 with the mirrored ellipses at both trait values (following equations 26) – (29). The values and stability of these ellipses were calculated using Mathematica. Only the upper diagonal graphs are shown. The white regions correspond to areas where no spatial variation in *N* or *z* was predicted in the system or where the predicted equilibrium was unstable or unreachable (negative *N*, or a nonzero imaginary part in *N*). Note that even in regions with stable density variation and or trait diversification it may depend on the initial conditions whether such an equilibrium is attained or not. The black dots correspond to the parameter settings that were used in the top panels of Fig. 7. In SI S2, the same figure, but including the unstable regions is shown, while in SI S1, a numerical verification of these results is shown.

Simultaneous trait diversification and variation in abundance exists and depends critically on the ratio between *c*_5_ and *c*_6_. As noted above, the selection gradient vanishes when *c*_5_ + *c*_6_*N*_1_ = 0. This happens at the critical density 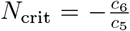. Above and below this threshold, selection acts in the opposite direction. If one of the two patches of a system is below *N*_crit_ and the other above, the trait value will diverge between the patches. In the example shown in Fig. 7, *N*_crit_ = 0.56. Given the sign of *c*_5_ (positive) and *c*_6_ (negative) that we used, evolution in patches with a density below 0.56 is towards higher values of *z*, while patches with a density above 0.56 tend towards lower values of *z*, as indicated by the grey line and color gradient in Fig. 7(d). However, if *c*_5_ and *c*_6_ would have the same sign, *N*_crit_ would be negative, and both patches will always have a density higher than *N*_crit_. In this case the direction of evolution is density-independent and only depends on the sign of *c*_5_ and *c*_6_. This is visible in the top right panel in figure 8, where all regions of stable trait variation lie in the quadrants where *c*_5_ and *c*_6_ have the opposite sign. When altering only *c*_5_ or *c*_6_, but not the other, stable coexistence can only occur when the sign of the coefficients does not change (the first five columns of the two top rows in Fig. 8). The importance of the ratio between *c*_5_ and *c*_6_ is further stressed in the topright panel in Fig. 8. If the regions with stable variation in both trait value and abundance would be determined by the effect of *c*_5_ and *c*_6_ on *N*_crit_ only, we would expect the region to be demarcated by two straight lines through the origin in the *c*_5_*, c*_6_-panel. However, the values of *c*_5_ and *c*_6_ not only affect *N*_crit_, but simultaneously the values of the equilibria, which is why the actual regions for stable variation in trait value and abundance deviate somewhat from the area between the imaginary straight lines through the origin.

Compared to the ecological model, *c*_5_ and *c*_6_ introduce a linear trait dependence to *c*_0_ and *c*_1_ respectively. In Fig. 8, this is visible in terms of a strong negative relation between *c*_0_ and *c*_5_ as well as *c*_1_ and *c*_6_ for cases where long-term variation in *N* can be maintained, visible as a turquoise diagonal band in the upper left quadrant of the *c*_0_*, c*_5_ and the *c*_1_*, c*_6_ panels. Similarly, also for the region where variation in both *z* and *N* can be maintained, larger values of *c*_1_ allow for more negative values of *c*_6_.

As for the lower four rows in the graph, the regions where variation in abundance can occur together with trait diversification, are generally a subset of the regions in Fig. 5 where variation in abundance was long-term stable. The values for *c*_0_–*c*_4_ that were used to produce the parameter space figure of the evolutionary model were identical to those used for the parameter space figure of the ecological model. If *z* were kept to 0 in the evolutionary model, we would retain the ecological model. However, visually, from Fig. 5 and 8 it seems that the possibility of the trait to evolve to a value of 1, in our specific model, seems to largely divide the regions where stable variation in abundance can occur into those where this can happen together with trait diversification and those without, without changing the general shape of these regions. Furthermore, for our focal parameter combination at least, inclusion of trait evolution produced a few novel regions where stable variation in *N* can be maintained.

### 2.4. Migration

To explore the robustness of our results to migration between patches, we included migration terms into both the ecological and eco-evolutionary model. We assume that each individual migrates to the respective other patch with a rate *m*, independently of the current population density or the individual’s trait value. Detailed methods and results are described in SI S4. In brief, we find that our results on the emergence of spatial density variation and trait diversification are robust to small amounts of migration between patches. With increasing migration rate, spatial heterogeneity decreases and eventually breaks down, first for traits and then for densities. In the example in Fig. 9, the smallest non-zero migration rate leads to both spatial density variation and trait variation, as in the model without migration. An intermediate migration rate still allows for spatial density variation, but trait variation disappears. And for the highest migration rate, both population densities and traits become homogeneous in space.

**Figure 9:**
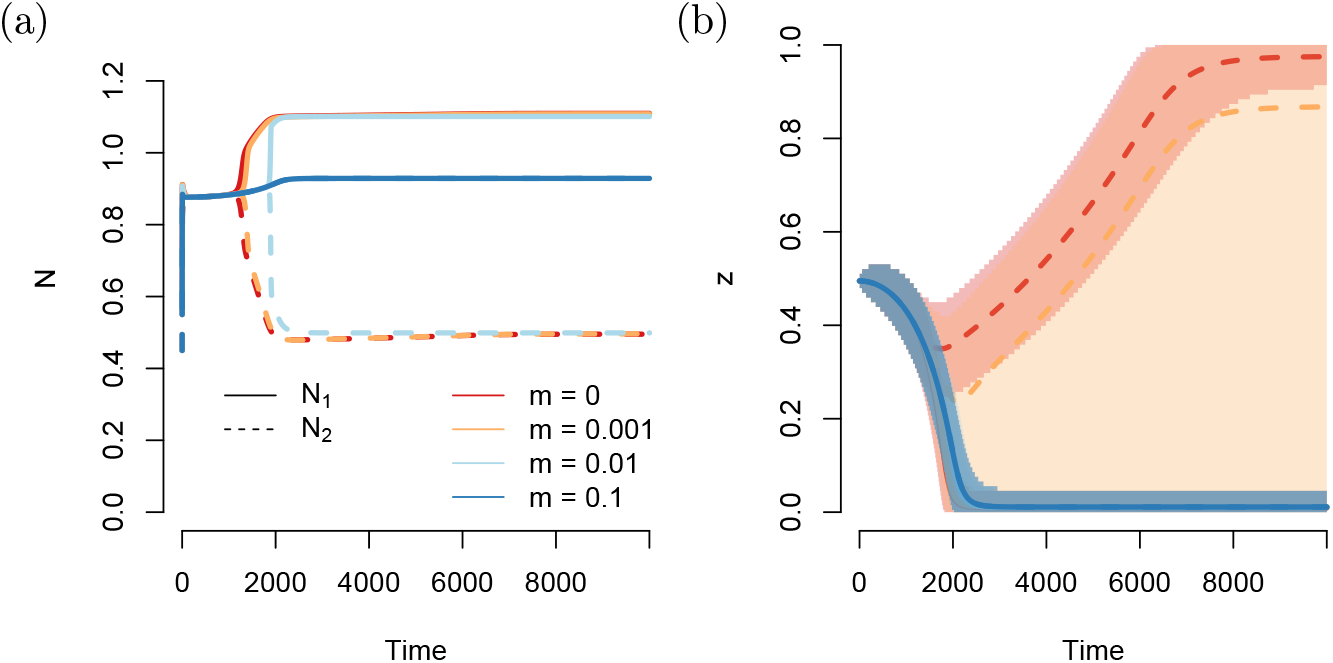
Time series from numerical solutions of the eco-evolutionary model with migration (see S4 for details) and an initial trait value *z*_0_ = 0.5. The other parameter values are the same as in Fig. 7. The shaded regions in panel (b) show the region in trait space that contains 95% of the population of each patch. For migration rates 0.01 and 0.1, both patches reach the same final average trait value; these lines completely overlap. When migration is absent, a high average trait value is reached in the low density patch. For *m* = 0.001, a high average trait value is still achieved in the low density patch, although the large shaded region implies that the patch also contains a non-negligible fraction of low-trait value individuals. With increasing migration rates, first the adaptation disappears (light blue lines and shaded regions overlapping with the blue line and region in panel b) and when *m* = 0.1 even the difference in abundance between the patches disappears.

### 2.5. Individual-based models

In order to test how the results change in the presence of multi-locus genetics, as well as demographic stochasticity, we repeated parts of the analysis with an individual-based model (IBM, see SI S5 for details). With appropriate parameter choices, spatial density variation and trait diversification did occur in the individual-based model. An additional parameter in the IBM was patch area. With small patch area, patches could accommodate few individuals and demographic stochasticity was strong, leading to frequent extinctions. With large patch area, the number of individuals was larger in total, there was less demographic stochasticity and the results were more similar compared to the deterministic model.

## 3. Discussion

In this study, we have explored the ecological and eco-evolutionary consequences of multi-scale density dependence where an individual’s fitness depends not just on population density in its own patch but also on the density in another patch in the region. We have shown that multi-scale density dependence may lead to the emergence and stable maintenance of spatial variation in population densities in an otherwise homogeneous environment and to the diversification of traits under density-dependent selection. That is, without any extrinsic heterogeneity, different niches emerged in the population, with some individuals being better adapted to low-density situations and others to higher density. We also observed how spatial density patterns and trait variation influenced each other through eco-evolutionary feedbacks. Specifically, we have shown a case where spatial density variation arose only when traits were allowed to evolve. Our model thus emphasizes how eco-evolutionary feedbacks can qualitatively affect both trait and population dynamics.

### 3.1. Formation of density patterns in space

In nature, spatial variation in population density between patches or subpopulations of the same species is ubiquitous. Our study highlights one possible mechanism that can produce or contribute to spatial density variation. A variety of other mechanisms exist. Firstly, evidence from natural populations suggests, that abiotic or biotic environmental conditions, such as temperature, climatic stability, precipitation, or food availability are key drivers of variation in density (Santini et al., 2018). Observed spatial variation in density can, however, also simply be caused by stochasticity, although such variation will not be stable over longer time scales. Finally, there are explanations that require neither extrinsic heterogeneity, nor stochasticity. Our study falls in this last category.

Many models from this last category (e.g. Bolker and Pacala, 1997; Bolker, 2003; Sasaki, 1997) focus on the interplay between random, undirected dispersal in some local neighbourhood, and intraspecific competition with other individuals in some competition neighbourhood. If typical dispersal distances are small relative to the spatial scale of competition, clusters of high population density can form. Similar to our model, these models also require some degree of non-local competition for pattern formation. Stable density variation can also emerge as the product of dispersal directed towards high population density (see e.g. Ellis et al., 2019), the interplay between small-range facilitation and long-range resource competition (van de Koppel et al., 2005), interactions with other species, such as reproductive interference (Ruokolainen and Hanski, 2016), hostparasitoid or -parasite interactions (Boots and Sasaki, 2000; Hassell et al., 1994), and interspecific competition acting over a smaller spatial scale than intraspecific competition (Murrell and Law, 2003).

Other studies have considered Allee effects as a key ingredient for the emergence and maintenance of spatial heterogeneity. For example, Gyllenberg and Hemminki (1999) showed how Allee effects caused by mate-finding difficulties together with non-local competition can lead to stable density differences between patches, even with nonzero migration. Their model is similar to our ecological model, except we did not explicitly model a specific Allee effect. Another study has shown how Allee effects can cause a population to be completely absent from some areas while being present in others, thereby limiting the spread of an invasive species (Keitt et al., 2001). Spatial heterogeneity in the sense of presence-absence variation can also be explained by metapopulation models (as first developed by Levins, 1969), but since each patch in a metapopulation experiences recurrent extinction and recolonization events, density patterns will not be stable over time.

More generally, all population models with alternative stable states can generate stable spatial heterogeneity in population density under appropriate initial conditions and with sufficiently small migration between locations. But without feedbacks between the patches, i.e. multi-scale density dependence, symmetric situations would always be as stable as asymmetric situations. Our model allows for situations where asymmetric densities are the only stable equilibria at which the species can exist (see for example Fig. 4 top, Fig. 6).

### 3.2. Maintenance of trait variation and relation to other coexistence mechanisms

In our eco-evolutionary model, a necessary but not sufficient condition for the maintenance of trait variation is the simultaneous maintenance of spatial variation in population density (see Fig. 8). Under appropriate parameter combinations, stable trait variation emerges with a high-density specialist dominating in the high-density patch and a low-density specialist dominating in the low-density patch, essentially a case of local adaptation (see Fig. 7). At equilibrium, there is still a very small amount of within-patch trait variation around the optimal trait value due to mutation-selection balance.

Chesson’s coexistence theory is a powerful framework to understand and classify coexistence mechanisms under spatio-temporal heterogeneity (Chesson, 2000). Yet, the coexistence between a high-density specialist and a low-density specialist in our model does not appear to fit straightforwardly into this framework. Because we have two types and two limiting factors, e.g. the densities in the two patches or functions of them, we do not seem have one of the cases where invasion growth rates can be cleanly partitioned into contributions from fluctuation independent frequency-dependence, storage effects, relative nonlinearity etc. (Barabás et al., 2018). Instead, we here provide an intuitive reasoning for how mutual invasibility and coexistence are achieved in our model.

Mutual invasibility and therefore stable coexistence of a high-density specialist and a low-density specialist can be achieved in two ways in our eco-evolutionary model. The first scenario is that each strategy produces spatial variation in density on its own and the high-density specialist can invade the high-density patch of the low-density specialist and the low-density specialist can invade the low-density patch of the high-density specialist. This scenario is illustrated by the example in Fig. S3.1. The second scenario is that the low-density specialist on its own has the same density in both patches, a density that allows the high-density specialist to invade, and the high-density specialist produces spatial variation in density such that the low-density specialist can invade the less dense patch. This is the scenario in Fig. 7. If neither the high-density specialist nor the low-density specialist have spatial variation in density on their own, mutual invasibility does not appear possible in our model.

To our knowledge, the role of multi-scale density-dependent selection in the maintenance of polymorphism has not been investigated before. Engen and Sæther (2019), however, showed that density-dependent selection can affect spatial trait patterns in a model with dispersal and temporal environmental heterogeneity. Also, there is previous work on how the spatial scale of density regulation in patchy landscapes affects the maintenance of polymorphism (Ravigné et al., 2004). If density regulation happens locally after selection, this is called soft selection (Levene model, Levene, 1953). If density regulation happens globally, this is called hard selection (Dempster model, Dempster, 1955). In soft-selection models, population densities are usually not affected by selection or migration, whereas in hard-selection models they may be affected (Lenormand, 2002). It has been shown that compared to hard selection, soft selection is more conducive to maintenance of polymorphism in response to environmental heterogeneity (e.g. Ravigné et al., 2004). More recently, also mixtures of hard and soft selection have received attention (De Meeûs and Goudet, 2000; Débarre and Gandon, 2011). The environmental heterogeneity in these studies, however, was assumed to be unrelated to population density. In our model, selection and density-regulation cannot be clearly separated and thus our model does not fit perfectly into the hard-selection/soft-selection framework. It shares more aspects with hard-selection models, most importantly that fitness influences absolute number of offspring and population densities, but there are also many additional aspects in our model like Allee effects and feedbacks between density and selection.

### 3.3. Migration and stochasticity

As described in section 3.1 and SI S4, migration (and dispersal) can have profound effects on the maintenance of variation in abundance. Like our ecological model, our evolutionary model is also robust against small amounts of migration (Fig. 9 and SI S4). However, with increasing migration rates, the accompanying gene flow between the patches hampers adaptation through gene swamping (Lenormand, 2002). This effect is expected to be particularly strong when the difference in abundance between the two patches is large, and a disproportionately large number of individuals migrate from the larger to the smaller patch. In a system where the patches have more similar abundances, gene swamping is thus expected to have a smaller effect, although the evolution of density-dependent traits would also be slower due to a weaker selection gradient. At higher migration rates, migration hampers not only adaptation, but even leads to the system reaching an equilibrium in which both patches have the same abundance. We thus only see negative effects of migration on long-term differences in abundances and trait values between the two patches in the deterministic models.

Several pathways exist through which migration (or dispersal) can have positive effects on local adaptation. Specifically for evolution, trait-dependent migration can lead to trait diversification. Homogeneous migration may also aid adaptation, for example by resupplying alleles that have been lost to drift (Blanquart et al., 2012), or by preventing a patch from going extinct, leading to more time for an adaptation to spread (Gomulkiewicz et al., 1999). In our deterministic model, these effects do not play a role because migration is unstructured, and alleles can always re-emerge. When demographic stochasticity is included, as in our individual based model (SI S5), migration can have positive effects on diversification. In those model runs, migration counteracted stochastic extinctions of the smaller patch for long enough, such that adaptation to low density could take place, similar to the effect described by Gomulkiewicz et al. (1999). This effect disappears when the absolute number of individuals increases, thereby weakening the effect of demographic stochasticity on population level processes.

Stochasticity may also promote trait diversification regardless of migration. Due to the symmetry in the deterministic model, if the two patches have the exact same initial abundance and trait distribution, they cannot diverge over time. In such cases, small amounts of (demographic) stochasticity may lead to small differences between the patches that are then amplified by the internal model dynamics, leading to variation in abundance as well as trait diversification.

### 3.4. Interpretation and applications

Our model assumes that density dependence plays out on more than one spatial scale. Such multi-scale density dependence should be common because an individual’s fitness depends on many different processes and factors, such as juvenile survival, protection from disturbances, resource competition, mate finding, competition for mating partners, reproduction, interactions with other species etc., which will generally occur over different spatial scales (see e.g., Cook et al., 2001; Gascoigne et al., 2005; Rietkerk, 2004). However, not all forms of multi-scale density dependence will lead to spatial variation in population densities and trait diversification. In our model, when there is positive density dependence at low density at both scales, spatial density variation can only be stable if the positive effects of conspecifics at low density and the negative effects at high density are either both strongest for the own patch or both strongest for the other patch. However, even then, not all parameter choices lead to stable density variation (compare Fig. 4(a) to (c)). Moreover, in almost all our examples with stable variation *c*_1_ < *c*_2_ with a positive *c*_2_ and *c*_1_ being either negative or positive, suggesting that facilitation from the other patch is more important for density and trait variation than facilitation by individuals in the same patch. However, we can currently not generalize these claims for all of parameter space.

The requirements for stable density variation and trait diversification could be fulfilled for example in plant-pollinator systems where plants grow in two patches but pollinators are more mobile and can visit both patches. A focal plant patch may benefit from a small nearby patch, by guiding pollinators to the focal patch. However, when density in the nearby patch gets too large, pollinators may instead choose to spend most of their time at that nearby patch. Simultaneously, within the focal patch, high density may lead to higher resource competition, while low densities may make the patch difficult to find for pollinators. When these processes lead to asymmetric abundances across the patches, this in turn affects the optimal investment that plants should make in competitive ability. This long-term difference can then lead to the emergence of trait variation. Evolutionary processes can also affect the abundances of the system, and can lead to a shift from a symmetric to an asymmetric equilibrium in abundances and subsequently to adaptation in density.

The patches in our model can also be interpreted in terms of social groups rather than locations, such as bark beetles attacking a tree (Raffa et al., 2008), or cooperative breeders. Similarly, the two groups can also be interpreted in terms of two competing (identical) species in a single patch, as modelled by Gerla and Mooij (2014). Although we had originally not thought of this interpretation, it turns out that mathematically, the model described by Gerla and Mooij (2014) is nearly equivalent to the ecological version of our model without evolution and migration. They find the same equilibria and shapes of the isoclines, but do not fully assess the stability of the asymmetric equilibria. Interestingly, Gerla and Mooij (2014) also mention plant-pollinator dynamics as a biological example, noting specifically the similarities between their system and the plant-pollinator model by Lutscher and Iljon (2013). A similar interpretation could be applied to our study as noted above. However, our study differs by focusing on spatial dynamics, deriving a direct equation for the stability of the unstable equilibrium, and by also evaluating the effects of trait evolution, migration and stochasticity.

The actual occurrence of the here-described eco-evolutionary effects require empirical evidence. It is currently unclear whether the sets of coefficients that allow for spatial density variation and trait diversification occur in natural populations. The fitness coefficients could empirically be estimated by evaluating how the fitness in one patch varies when its density and the density of a nearby patch are manipulated. Alternatively, one could experimentally test predictions of our model. In our model, if an asymmetric equilibrium with one patch at high density and one patch at low density is stable, the mirrored situation with the respective other patch at high or low density should also be stable. Our model thus predicts that if one manipulated a system with asymmetric densities, for example by adding individuals to one patch and removing individuals from the other patch, the system should eventually shift from the basin of attraction of one asymmetric equilibrium to the other asymmetric equilibrium and stay there. By contrast, if the spatial density variation is due to extrinsic environmental differences, the system should return to the original equilibrium. Once it has been established that the system is ecologically capable of reaching asymmetric abundances, experimental evolution may be attempted. Although such experiments are challenging, testing the effects of eco-evolutionary dynamics can and has been done, for example by frequently replacing individuals in an 30 experimental population by wildtype individuals from a stock population, thereby disabling the evolutionary part of the feedback loop and comparing such a population to a population where evolution is allowed to take place (e.g. Hart et al., 2019).

### 3.5. Limitations and future work

We assumed a homogeneous environment, which may be unrealistic for most natural populations. Whether the effects described here are of importance when the environment is heterogeneous remains to be investigated. This could be evaluated through a model that also incorporates environmental heterogeneity for example through an additional term in the fitness function. Depending on the spatio-temporal pattern of environmental variation, many different model behaviors may be possible. We expect, however, that our results will be robust to some degree of environmental heterogeneity. Even if the patches are slightly different in their properties, the eco-evolutionary feedbacks described here should still act and while it might be more likely for the patch with slightly better environmental conditions to become the high density patch, depending on the starting conditions it may also be the other way around if the eco-evolutionary feedbacks are strong enough. The phenomena described here could also amplify the effects of environmental heterogeneity on spatial density variation. This interplay between intrinsic and extrinsic factors for the maintenance of eco-evolutionary variation is a promising direction for future research.

Temporally strongly varying environments may favor plastic rather than evolutionary responses: phenotypic plasticity could allow an individual to cope well with the different environments that it will experience over its life time. In our model, the environments that individuals and their offspring encounter critically depend on the migration rate and we would expect phenotypic plasticity to be favorable when the migration rate is small enough for spatial density variation to persist, but too large for evolutionary trait diversification. Generally, whether plasticity can evolve depends on the costs and limits of the plasticity (DeWitt et al., 1998), such as the degree of unpredictability of the future environment (Reed et al., 2010), in conjunction with the migration rate and the accuracy of the plastic response (Sultan and Spencer, 2002). The precise conditions should be explored in future extensions of our model.

Another potential key factor that determines the model behaviour is the spatial structure. We have investigated a model with a particular, very simple spatial set-up: two patches that mutually affect each other. An important direction for future work is to investigate how the eco-evolutionary dynamics explored in this study play out in other spatial settings, for example landscapes with more than two patches, or continuous space. In such scenarios, individuals could respond to spatial density variation in multiple ways, e.g. to the density in their own patch or neighborhood and to the density in the rest of the landscape, or separately to the densities in each patch, or as defined by a kernel based on distance. In continuous-space models, local dispersal is another mechanism that could interact with multi-scale density dependence to promote spatial variation in density. We speculate that at larger scales too, multi-scale density dependence may promote density and trait variation, although this needs to be confirmed with additional models.

Individuals in our model differ in just one trait and the fitness effects of this trait depend on just the population density in the individual’s own patch, even though fitness itself is affected by density on multiple scales. Future models could incorporate traits that are involved in density-dependent processes at a larger scale and whose fitness effects therefore depend on the density in the other patch or on total density. We speculate that similar eco-evolutionary feedbacks as we reported here would act in this case, which should also allow for the maintenance of trait variation under appropriate conditions.

Although much remains to be investigated, we argue that multi-scale density dependence is a common, potentially very important phenomenon in evolutionary ecology. Based on many empirical examples (see e.g., Cook et al., 2001; Gascoigne et al., 2005; Rietkerk, 2004) and the argument that fitness depends on multiple biological processes that will generally not play out at exactly the same spatial scale, we expect that multi-scale density dependence will be the rule rather than the exception. More generally, when studying natural systems, the outcome may depend on the, sometimes arbitrarily, chosen spatial scale over which the study is conducted (Kareiva, 1990; Murphy, 1989; Ray and Hastings, 1996; Snyder and Chesson, 2004) and multi-scale density dependence could contribute to explaining some of these inconsistent results. Finding relevant literature and synthesizing information on multi-scale density dependence is, however, made difficult by the lack of clear terminology for this phenomenon. Because of its important ecological and eco-evolutionary consequences which we highlighted in this study, we argue that multi-scale density dependence should receive more concentrated research attention.

## 4. Conclusion

Our study shows how in two mutually affecting patches, long-term abundance differences can emerge, even in the absence of external differences between the patches. Furthermore, we have shown how selection can lead to phenotypic diversification in traits. Because of eco-evolutionary feedbacks, the inclusion of mutations can lead to both ecologically and phenotypically different outcomes. Our study serves as a proof-of-concept, which we hope will inspire empirical validation and contribute to our understanding of possible mechanisms through which spatial variation in density and traits may emerge.

## 5. Acknowledgements

This research was funded by the German Research Foundation (DFG) as part of the SFB TRR 212 (NC^3^). Furthermore we are grateful to Hannah Živković and Matthias Kubacki for their contributions to the individual based model.

## 6. Author contributions

KvB and MW conceived the ideas and designed the methodology. KvB led the analyses and the writing of the manuscript. MW contributed extensively to the writing and interpretation of the results.

## 7. Supporting information

### S1. Numerical verification of the parameter space of the evolutionary model

The results in Fig. 8 were based on the Jacobian and the equilibrium values returned by Mathematica. However, these results are subject to the numerical precision of the computer, which is why we assumed imaginary parts of the equilibria smaller than 10^−16^ to be equal to zero and hence as reachable equilibria. We therefore numerically verified the results from Fig. 8 using the package deSolve (Soetaert et al., 2010).

### S1.1. Methods

Because the numerical solutions run slower compared to the analytical results, we performed these at a lower resolution, by dividing the coefficient range (−2 – +2) in 20 steps only. Hence every subpanel consists of 20 × 20 pixels (and checks). For every parameter setting (i.e. pixel), we first calculated the six possible nonzero equilibria that allow for variation in at least either trait value or abundance, using the analytical solution. These were the four possible mutual intersections of the ellipse corresponding to trait value 0 and trait value 1, and the asymmetric equilibria for a system where trait values were the same in both patches (either both at 1 or both at 0, considering only one of the mirrored asymmetric equilibria in each case). Then, we removed the imaginary parts of these results. Furthermore, we remove the equilibria with a negative real part, since these are not biologically relevant.

Subsequently, we evaluated whether the remaining equilibria were stable. For each of the equilibria 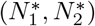, we pick four possible starting densities, that differ from the expected equilibrium by 2.5%. Each patch can be either 2.5% higher or lower, leaving us with the four possible starting conditions: 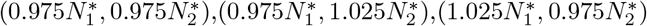, and 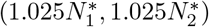. We call an equilibrium stable with respect to the density, if for each starting condition, after 100 time steps the difference between the abundance in the patches and their expected value according to the equilibrium is smaller than 2.5%.

In order to make sure that the population goes to the expected equilibrium, we had to set *μ* = 0, because when mutations are present, the trait values will deviate from 0 or 1 by a small amount, due to the mutation-drift balance and this deviation in *z* may subsequently lead to a small difference in abundances as well. Thereby hindering a direct comparison between the numerically predicted density and the expected density. However, this means that we have to assess the stability of the equilibrium with respect to *z* separately. We did so, by changing the initial trait distribution. Instead of putting the full population in either the bin with *z* = 0 or *z* = 1, we put 99% of the population in the extreme bin and 1% in the adjacent bin. When *z* = 0, this corresponds to:

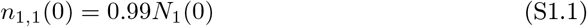

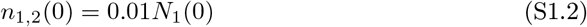

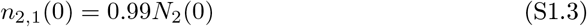

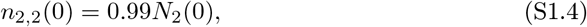

and zero density in all other bins. When *z* = 1, bins 100 and 99 take the role of bin 1 and 2 respectively. If the relative number of individuals in the extreme bin had increased at *t* = 100 compared to the initial value, the system was considered to be stable with respect to trait value.

### S1.2. Results

The resulting figure (Fig. S1.1) confirms the findings from Fig. 8.

**Figure S1.1:**
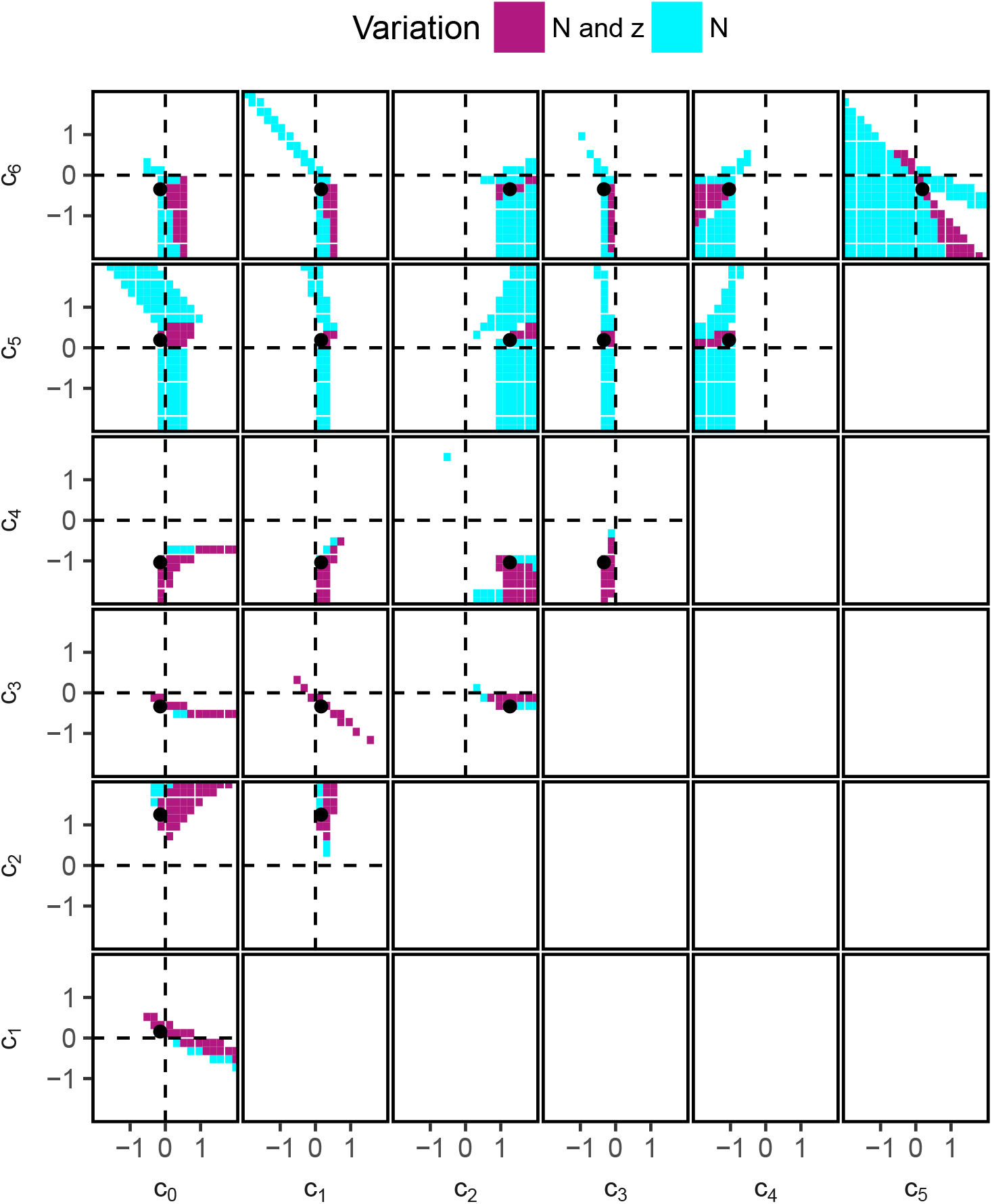
Regions in parameter space where spatial variation abundance and diversification of trait values can occur. The shown regions are based on numerically evaluating the population dynamics around the the asymmetric, equilibria that were obtained by intersecting the ellipses with trait value *z* = 0 and *z* = 1 with the mirrored ellipses at both trait values (following equations 26) – (29). The white regions correspond to areas where no spatial variation in *N* or *z* was predicted in the system or where the predicted equilibrium was unstable or unreachable (negative *N*, or a nonzero imaginary part in *N*). The black dots correspond to the parameter settings that were used in the top panels of Fig. 7. The results in thsi graph numerically confirm the findings from Fig. 8.

### S2. Parameter space of the eco-evolutionary model including unstable equilibria

**Figure S2.1:**
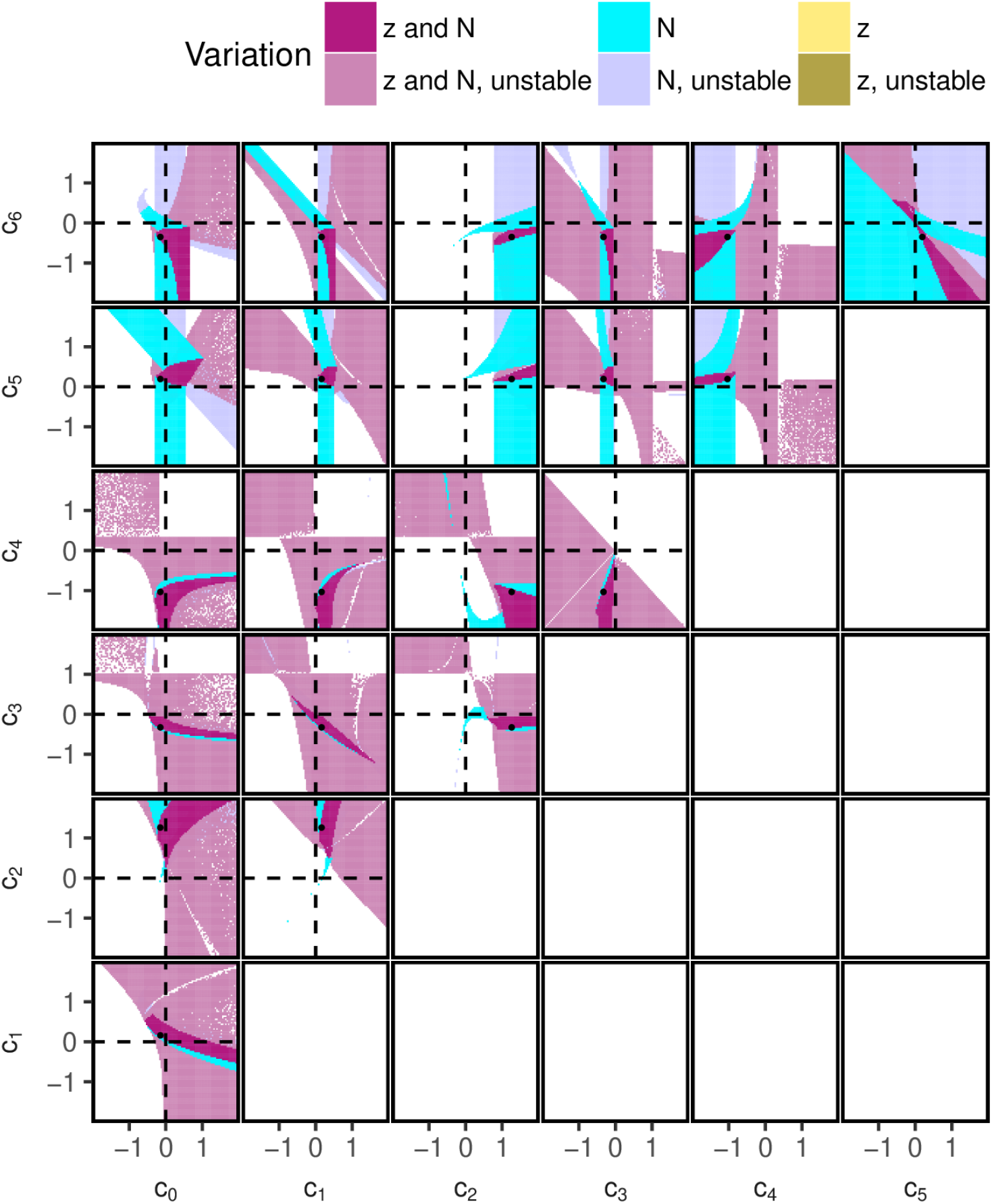
Regions in parameter space where spatial variation abundance and diversification of trait values can occur, including unstable equilibria. The shown regions are based on the properties of the asymmetric equilibria obtained by intersecting the ellipses with trait value *z* = 0 and *z* = 1 with the mirrored ellipses at both trait values (following equations 26) – (29). The values and stability of these ellipses were calculated using Mathematica. Only the upper diagonal graphs are shown. The white regions correspond to areas where no spatial variation in *N* or *z* was predicted in the system or where the predicted equilibrium contained negative values. The black dots correspond to the parameter settings that were used in the top panels of Fig. 7. The pixelated regions are due to the floating point precision in calculating the imaginary part.

### S3. Additional example for the eco-evolutionary model

Here, we show an additional example time series of the eco-evolutionary model. In this example, we have weakened selection by dividing both *c*_5_ and *c*_6_ by 10. In order to speed up selection in this example, we have increased the mutation rate *μ* to 0.05. All other parameters are as in Fig. 7. In this example, at both *z* = 0 as well as *z* = 1, the model has two equilibria, of which one is above *N*_crit_ and the other below.

**Figure S3.1:**
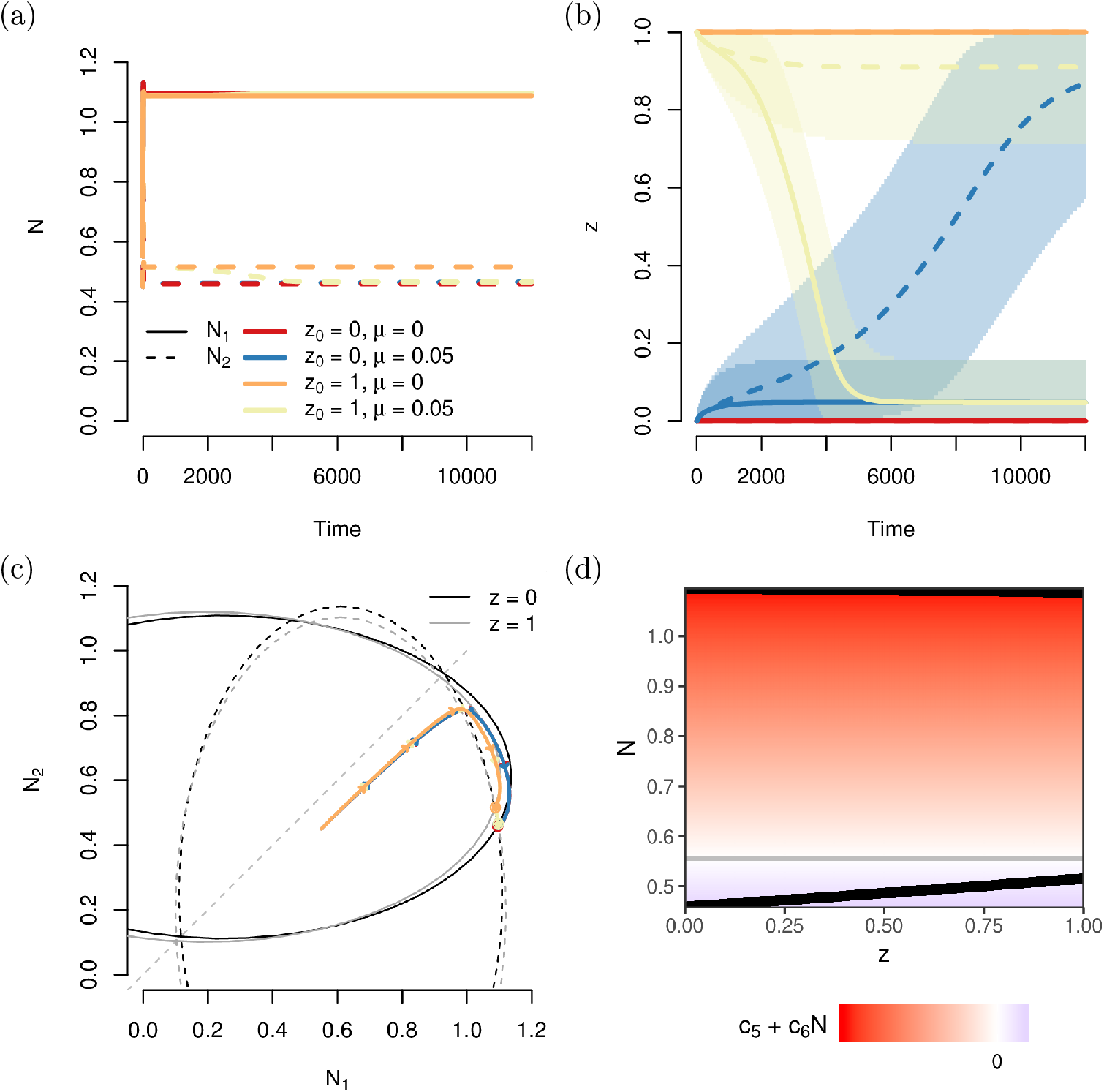
Example time series for the eco-evolutionary model. Different colors represent different model runs where mutations were either present (*μ* = 0.05) or not (*μ* = 0) and that started with a monomorphic population with either trait value 0 (*z*_0_ = 0) or 1 (*z*_0_ = 1). (a) Abundance time series, the red and blue trajectories largely overlap for the first 2000 time steps, as do the yellow and orange lines. (b) Corresponding mean trait values and the spread, shown as the regions in trait space that contained 95% of the individuals of each patch. For the scenarios without mutations (orange and red line), the trait values in both patches completely overlap. For the scenarios with mutations the trait values in the two patches diverge over time, regardless of the initial trait distribution. (c) Trajectories in phase space. Also drawn are the isoclines at *z* = 0 (black) and *z* = 1 (grey). (d) Equilibrium densities for monomorphic populations with trait value *z*. At any given value of *z* two branches exist, indicating the two different densities that the two patches will tend to. In the background, the direction of selection at any given density is shown, with red values referring to selection for smaller trait values and blue colors to selection for larger trait values. The grey line corresponds to *N*_crit_, the density at which selection vanishes. Parameter values: *c*_0_ = −0.148, *c*_1_ = 0.162, *c*_2_ = 1.262, *c*_3_ = −0.326, *c*_4_ = −1.034, *c*_5_ = 0.0194 and *c*_6_ = −0.03492.

### S4. Migration between patches

Differences in abundance between patches can be evened out through migration. If the same proportion of individuals in both patches migrates to the respective other patch, the larger population will contribute more individuals to the smaller population and vice versa. Hence, the difference in abundance between the patches is expected to decrease and the smaller patch should be less likely to go extinct. Furthermore, migration may hamper trait diversification through recurrent inflow of genes that have emerged through a selection pressure elsewhere. It is therefore vital to see whether the diversity in abundance and trait values that emerged in our original model can be maintained under migration.

#### Ecological model

First, we adapt the purely ecological model to include migration:

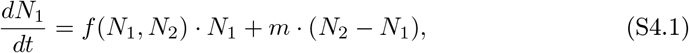

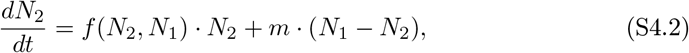

with *m* being the fraction of individuals in each patch that migrate to the other patch. The fitness function *f* is the same as the one used in the main text (equation 2).

**Figure S4.1:**
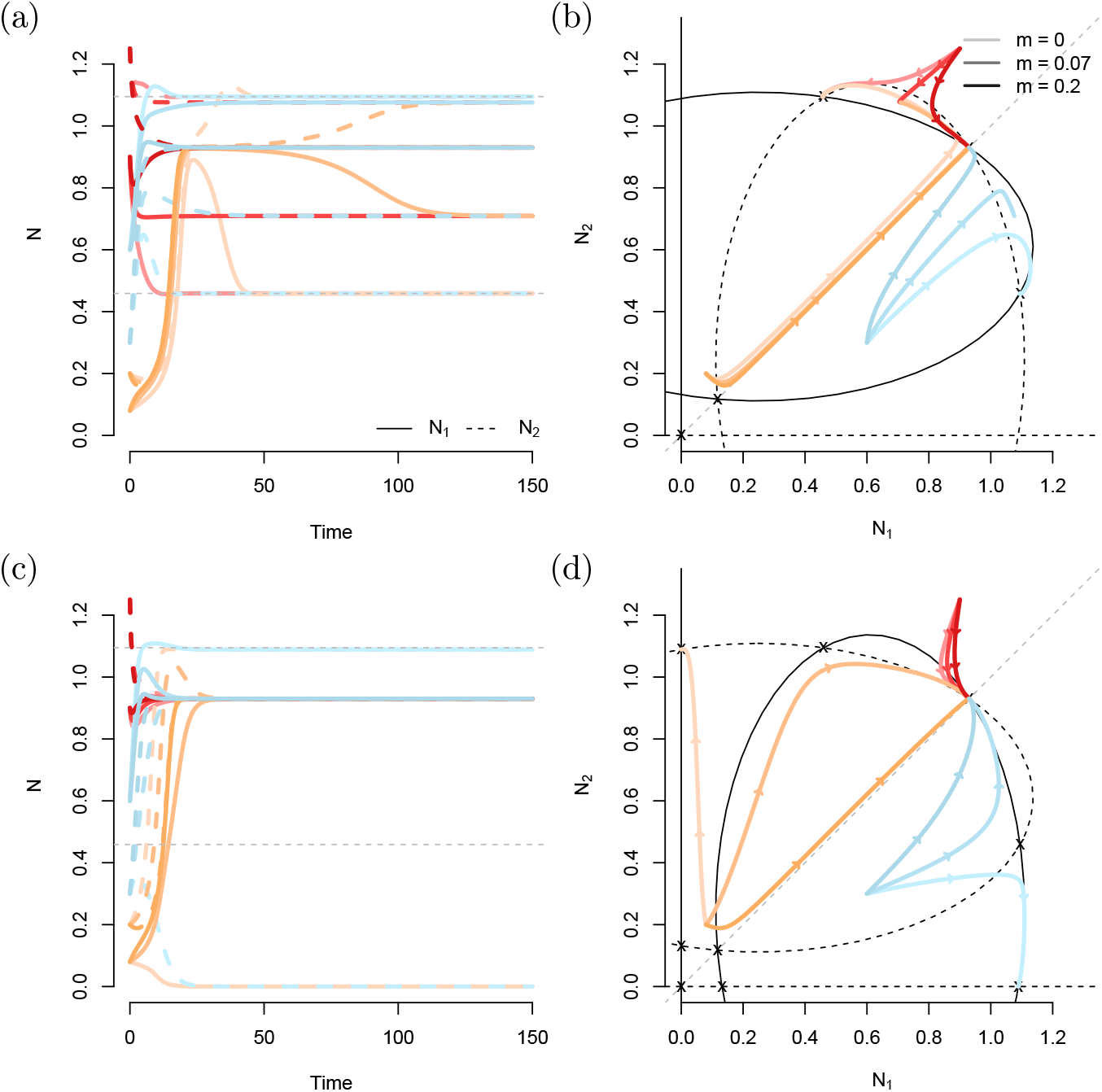
Time series of abundance for the ecological model with different migration rates. Otherwise, the parameter settings are as in the main text (see Fig. 4). The different colors correspond to different initial conditions, with the saturation indicating the migration rate. The equilibria and ellipses are shown for the case without migration.

We solved the model with migration numerically and compared the results to those without migration (Fig. S4.1). Migration increases the time it takes for the system to reach the asymmetric equilibrium (top left panel). Furthermore, migration affects the existence/stability and precise value of the equilibria (compare the red trajectories for *m* = 0 and *m* = 0.07 in the top panels). This becomes apparent when plotting the final abundances in both patches as a function of the migration rate (Fig. S4.2). At low migration rates, the two separate equilibria remain relevant. However, when the migration rate reaches a critical value (close to 0.10 for the depicted parameter combination), the system always tends to a symmetric equilibrium. Note how in this case the density in the smaller patches increases faster with the migration rate than it decreases in the larger patch. As a consequence, the total population size increases as a function of the migration rate. This is a well-known effect that is caused by us not explicitly modelling the underlying causes for the density dependence (Wang and DeAngelis, 2019).

**Figure S4.2:**
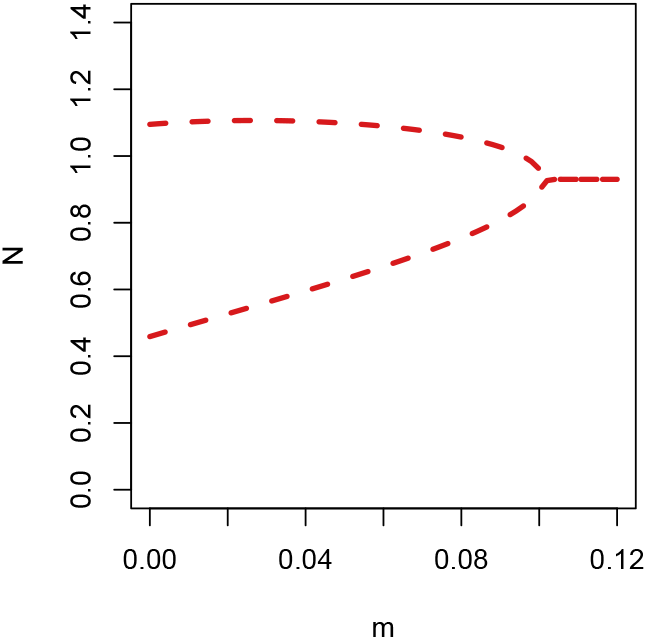
Final densities in the two patches in a numerical solution after the first 10^5^ time steps as a function of the migration rate. This was enough time for the system to reach the equilibrium. Results did not vary among the three different starting conditions from Figure S4.1.

#### Evolutionary model

In the evolutionary model, migration is included analogously; so for patch 1, the differential equations become:

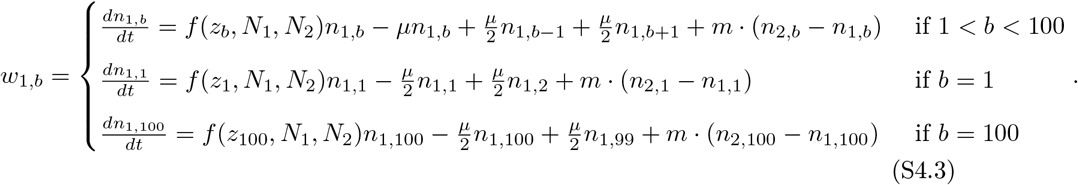

In our approach, we treat *m* as a constant, such that migration is trait-independent.

From numerical solutions of the evolutionary model with migration (Fig. 9), it becomes apparent that the difference in average trait value is far more sensitive to migration than the difference in abundance at the used parameter values: at *m* = 0.01, the two patches still reach different abundances, but they no longer obtain different average trait values. Therefore, we show the final abundances of time series that were solved numerically for up to 10^4^ time steps in two different graphs with different scales (Fig. S4.3). The differences in trait value disappear rapidly with increasing migration rates. This seems to be caused by the relatively large influx of individuals from the larger patch that are adapted to the density in their native patch, combined with a relatively low selection gradient. Interestingly, the effect of the migration rate depends on the mutation rate. This reflects that an increase in mutation rate affects the mutation-selection balance, causing the average trait value to shift away from the boundary values.

**Figure S4.3:**
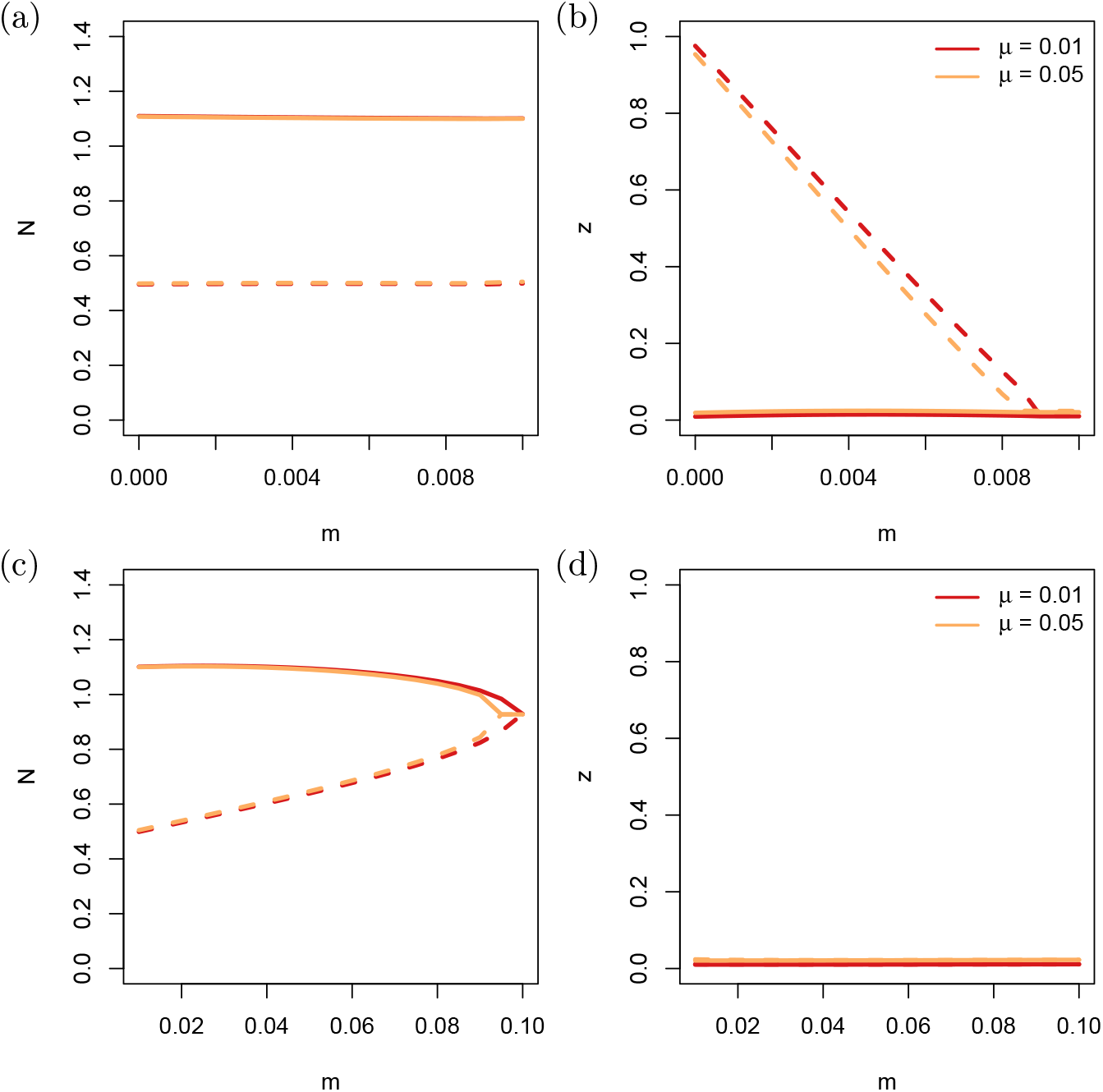
Final abundances in numerical solutions at time *t* = 10^5^, which was enough time for the system to reach equilibrium. The top and bottom half differ only in the range of migration rates that they contain. Mutation rate was varied, but all other parameter values were the same as in the evolutionary model in the main text.

In constructing Figure S4.3, we have re-assigned patch identity afterwards based on the final density (the patch with the larger final density was assigned to patch 1). Otherwise, the lines would move back and forth between the two branches. That does mean however, that the branches no longer correspond to a specific patch.

### S5. Individual-based simulations

We tested the robustness of the results against demographic stochasticity with an individual-based model. This also allowed us to include diploid multi-locus genetics.

#### Genetics

Genetics were diploid and consisted of a 10-locus system with 10 alleles per locus. Alleles were inherited independently across loci (no linkage). Every locus, contributed a value between 0 to 0.1. When adding all 10 loci together, this led to the total trait value ranging between 0 and 1. The value of a locus was determined by averaging the value of its two alleles. The alleles themselves had evenly spaced values between 0 and 0.1. There were no factors other than genetics that affected the trait value.

Finally, for every newborn, there was a probability *μ* that it obtained a mutation at one of its loci. If a mutation was determined to occur, the locus and the value of the target allele were drawn from a uniform distribution, independently of the original allele value.

#### Fitness function

We assumed non-overlapping generations and the fitness function only affected female reproduction. This is one of the deviations from our original model, where males and females were not distinguished. For a female in patch 1, with trait value *z*, the number of offspring was drawn from a Poisson distribution with mean:

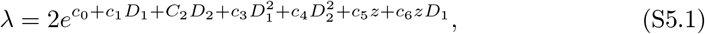

with *D_i_* the density in patch *i*. If patch *i* has area *A_i_*,

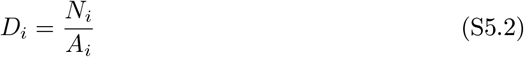

The factor of two in equation (S5.1) serves to compensate for the fact that males do not generate offspring. Instead, for every female that obtained offspring, the father of that clutch was randomly sampled among all males in the corresponding patch. Offspring were also born in the same patch, although in the model runs with migration, they had a small chance to move to the other patch. Although this process actually corresponds to dispersal, we use the term migration throughout this supporting information, for consistence with the main text.

Finally, we had to set an area for the patches. For all model runs, the area was equal in both patches (*A*_1_ = *A*_2_ = *A*). In the original model, the scaling of *N*_*i*_ was arbitrary, and *N*_*i*_ could also take non-integer values. In the individual-based model this is not possible. Here, *N*_*i*_ can only take integer values. Changing the size of the area allows us to alter the relative effect of demographic stochasticity, and thereby also the amount of genetic drift in the system. When the area is large, the same equilibrium density corresponds to a larger absolute number of individuals. Hence, demographic stochasticity is expected to have only a limited effect under these circumstances.

#### Initial population

The number of individuals in each patch was drawn from a normal distribution with mean *N*_0_ and standard deviation *σ* = 10. For each of the patches, the fraction of males in the initial population was drawn from a uniform distribution between 0.25 and 0.75. The number of individuals and the number of males were then rounded to the closest integer and applying the absolute value to avoid negative values (which rarely occurred because of sufficiently large *N*_0_). Next, the allele values at the loci were also randomly sampled from a uniform distribution: at every locus alleles were assigned, with equal probability for every possible allele.

#### Runs and results

Direct averaging of the replicates proved difficult, due to stochasticity making it impossible to predict which of the two patches would become the high density patch. Instead, we summarize the results qualitatively and show a few time series to illustrate the point. Throughout the simulation runs, we varied three parameters: *m* ∈ [0, 0.005], *μ* ∈ [0.1, 0.05], and *A* ∈ [100, 1000, 5000]. We tested all combinations of these parameter settings and ran 10 replicates for each combination, for a total 2 *·* 10^5^ time steps. If one of the two patches went extinct, the simulation was stopped.

**Figure S5.1:**
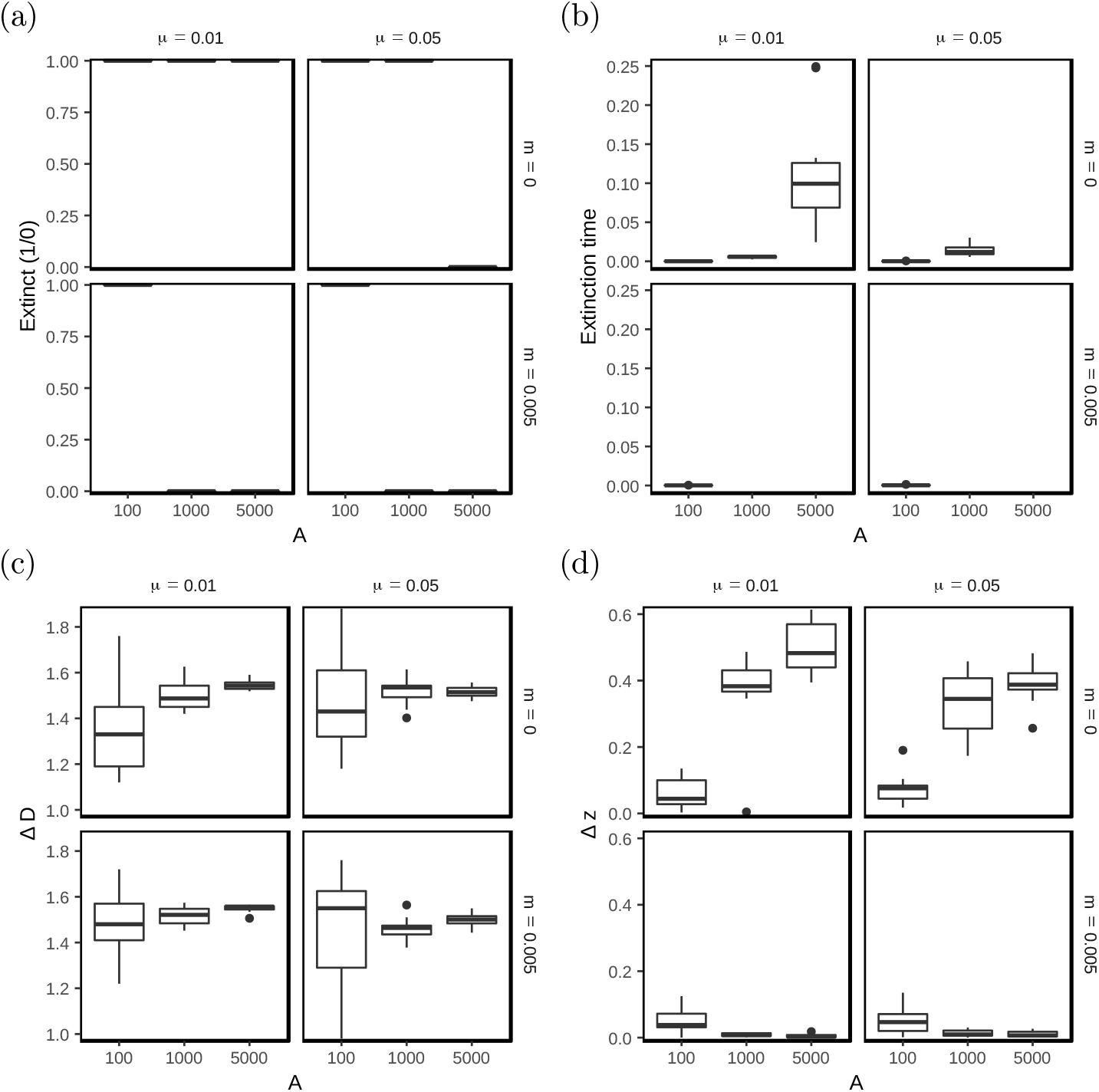
Summary of the runs from the individual-based model. Shown are boxplots that display (a) whether a patch in the replicates went extinct (1) or not (0), (b) the time at which this happened, (c) the difference in density between the patches at that point in time, and (d) the difference in average trait value between the patches at the end of the simulation (2 *·* 10^5^ or extinction time). The parameter values of the fitness function were as in Fig. 7.

First of all, the results show a strong consistency across replicates, despite the presence of demographic stochasticity. This stochasticity leads to extinction in all cases where *A* = 100 (Figure S5.1(a)). Furthermore, this panel also show that the mutation rate, and hence the potential speed of adaptation, has a direct effect on whether the populations can persist. Comparing panel (a) to (b), shows that at lower mutation rate, extinction occurs after approximately 10% of the simulation time, indicating that short-term co-existence of two patches with different abundances and trait value might be possible (as illustrated in Figure S5.2). Panel (a) also shows that extinction is less likely when migration is allowed. Migration has, however, only a very limited effect on the final difference in abundance (panel (d)). In agreement with the results from SI S4, migration strongly affects the possibility of trait adaptation (panel (d), and Figure S5.3). The situation that is closest to the deterministic evolutionary model that we presented in the main text consists of a large patch size that reduces the effect of demographic stochasticity, combined with low (no) migration and high mutation rates (for rapid adaptation). An example time series of this case is shown in Figure S5.4.

**Figure S5.2:**
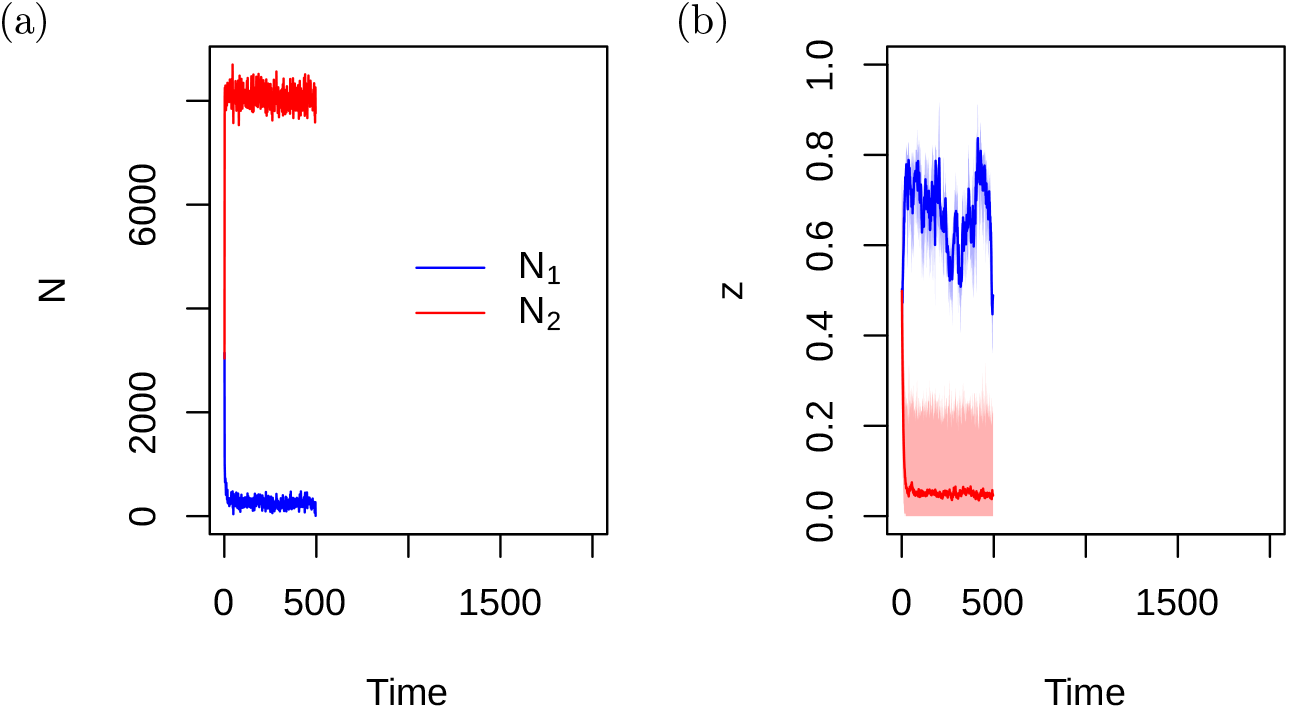
Time series run for one of the replicates with *μ* = 0.01, *m* = 0 and *A* = 5000.

**Figure S5.3:**
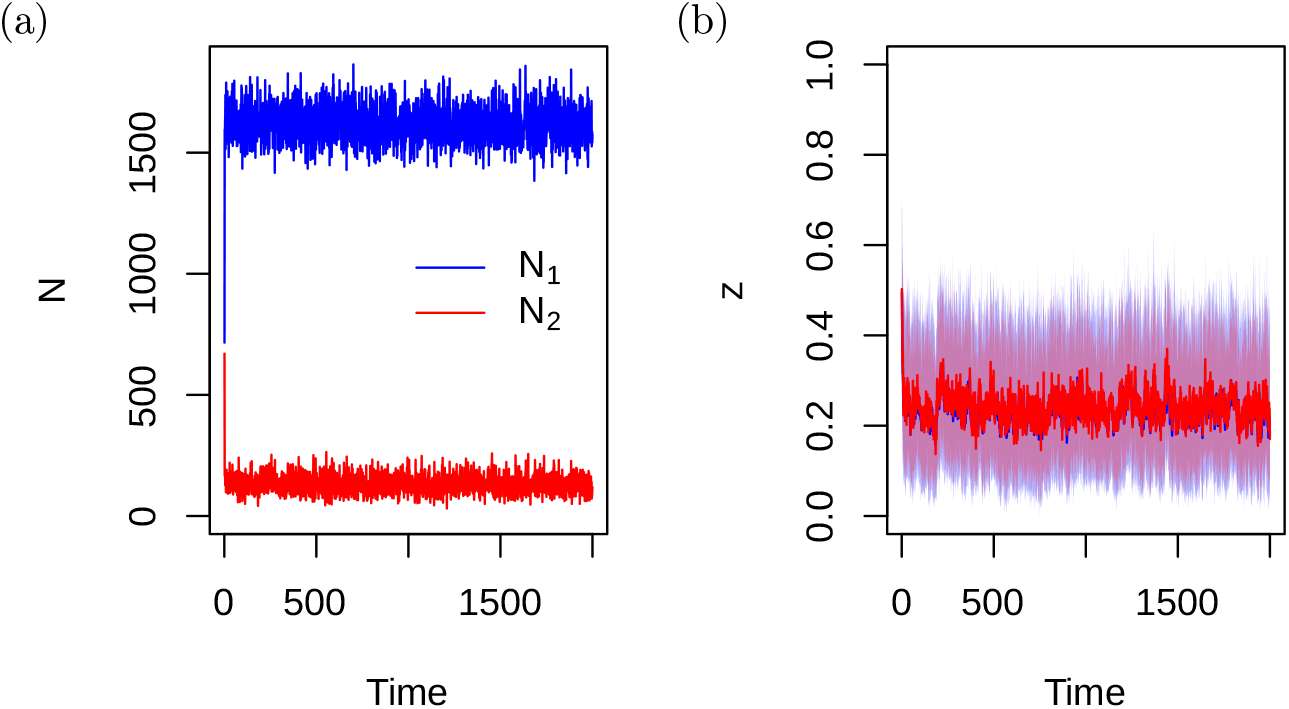
Time series run for one of the replicates with *μ* = 0.05, *m* = 0.005 and *A* = 1000.

**Figure S5.4:**
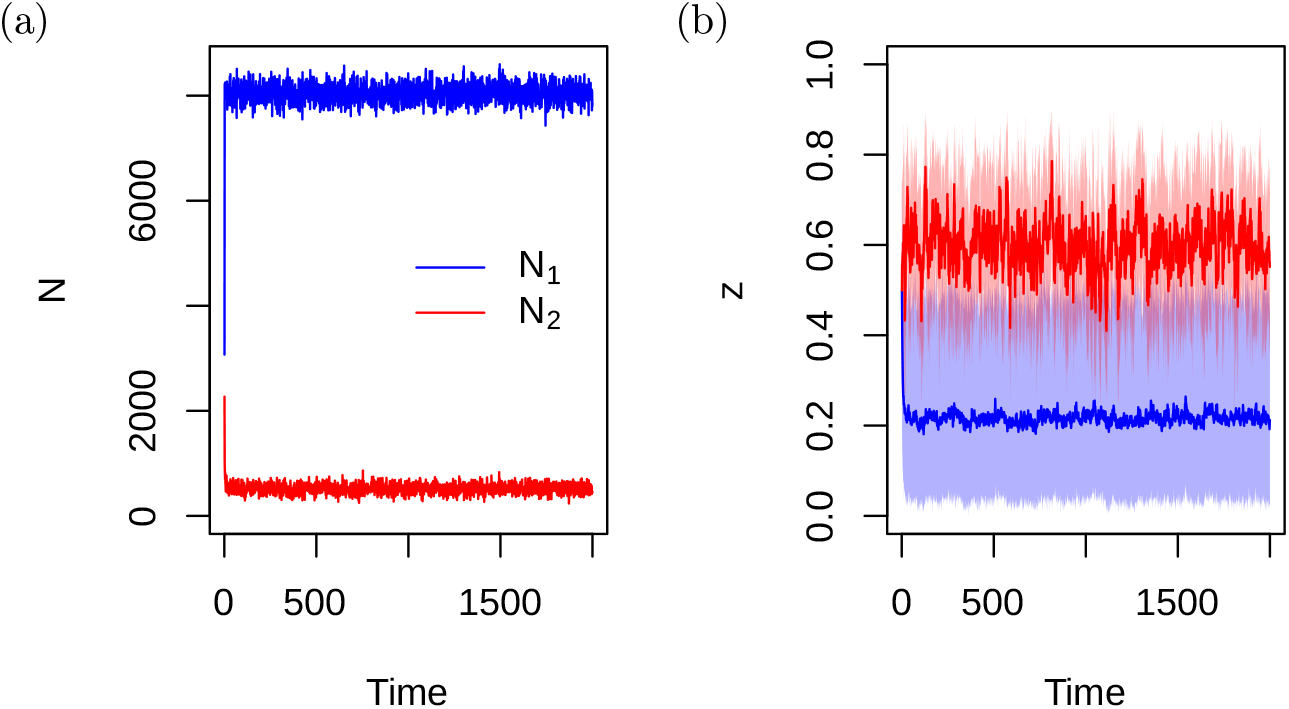
Time series run for one of the replicates with *μ* = 0.05, *m* = 0, and *A* = 5000.

